# Long-range chromatin interactions on the inactive X and at *Hox* clusters are regulated by the non-canonical SMC protein Smchd1

**DOI:** 10.1101/342212

**Authors:** Natasha Jansz, Andrew Keniry, Marie Trussart, Heidi Bildsoe, Tamara Beck, Ian D. Tonks, Arne W. Mould, Peter Hickey, Kelsey Breslin, Megan Iminitoff, Matthew E. Ritchie, Edwina McGlinn, Graham F. Kay, James M. Murphy, Marnie E. Blewitt

**Author notes:** Mammalian Development, Sir William Dunn School of Pathology, University of Oxford, South Parks Road, Oxford OX1 3RE, UK.

## Abstract

The regulation of higher order chromatin structure is complex and dynamic; however we do not yet understand the full suite of mechanisms governing architecture. Here we reveal the non-canonical SMC protein Smchd1 as a novel regulator of long-range chromatin interactions, and add it to the canon of epigenetic proteins required for *Hox* gene regulation. The effect of losing Smchd1-dependent chromatin interactions has varying outcomes dependent on chromatin context. At autosomal targets transcriptionally sensitive to Smchd1 deletion, we find increased short-range interactions and ectopic enhancer activation. By contrast, the inactive X chromosome is transcriptionally refractive to Smchd1 ablation, despite chromosome-wide increases in short-range interactions. There we observe spreading of H3K27me3 domains into regions not normally decorated by this mark. Together these data suggest Smchd1 has the capacity to insulate the chromatin, thereby limiting access to other chromatin modifying proteins.

---

In recent years there has been a dramatic increase in our understanding of how DNA forms long-range interactions^1-4^. This is largely due to chromatin conformation capture and sequencing (Hi-C) techniques, which have been used to map chromatin contacts genome-wide^5,6^. Hi-C revealed the presence of topologically associated domains (TADs): Mb-sized self-interacting regions^7^. As these techniques increased in resolution, they also enabled study of chromosome looping. However, our knowledge of these genomic structures has far preceded an understanding of mechanism and function.

As Hi-C data revealed the ways in which DNA molecules are spatially organised within the nucleus, models were developed to explain the mechanism by which chromatin structures could be formed and maintained. The model with the most traction has been the loop extrusion model, in which DNA extrudes through a ring formed by the SMC complex, Cohesin, until the complex is stabilised by convergent Ctcf binding at TAD boundaries. As such, Ctcf and Cohesin have been the focus of much research in the field. However, genome-wide loss of Ctcf and Cohesin mediated structures often have a surprisingly limited effect on transcription^8,9^. It is known that a host of epigenetic regulators are also involved in the formation and maintenance of higher order chromatin structures that modulate transcription. This is exemplified by polycomb repressive complex 1 (PRC1), which is involved in forming long-range repressive chromatin interactions that are distinct from TADs, and which result in compaction of the locus^10,11^. Continued identification of epigenetic regulators that mediate long-range chromatin interactions will help to resolve the function of many of the DNA structures defined using Hi-C based techniques and help to refine models that explain the principles of genome organisation.

Structural maintenance of chromosomes hinge domain containing 1 (Smchd1) is a non-canonical SMC protein^12^. Similar to the canonical SMC proteins^13-15^, Smchd1 possesses a hinge domain through which it dimerises and interacts with nucleic acids^16,17^, but differs in that the N-terminal ATPase domain belongs to the GHKL family, rather than the ABC-type ATPase domain present in canonical SMC proteins^18,19^. Smchd1 exhibits further divergence from the canonical SMC proteins; for example, no obligate Smchd1 binding partners have been identified, unlike the additional non-SMC component proteins that make up the Cohesin or Condensin SMC complexes. To date, the role of Smchd1 in regulating higher order chromatin organisation has not been studied, though work from our group has suggested Smchd1 may functionally oppose Ctcf^17^.

Smchd1 plays a role in epigenetic repression throughout the genome. We first identified Smchd1 based on its role in repeat-induced silencing in mice^12,20^, and it also plays an important role in silencing the *D4Z4* macrosatellite repeat in humans^21^. Smchd1 has a critical role in silencing the inactive X (Xi) chromosome in females^12,22^, and in transcriptional repression of specific autosomal loci, including select imprinted genes and the clustered protocadherins^17,23−^^25^. Each of these sites are direct targets of Smchd1^12,17,26^, but interestingly so are other autosomal loci including the *Hox* genes^17^, where the role of Smchd1 has not been explored.

The mode underlying Smchd1-mediated gene silencing is unknown, although several epigenetic alterations are seen at Smchd1 target loci in cells derived from *Smchd1* null embryos. *Smchd1* null females die around embryonic day 10.5 (E10.5), with failed X inactivation^12,22,23,25^. In these embryos the early stages of X inactivation proceed normally; however, by mid-gestation failed X-linked gene silencing is accompanied by DNA hypomethylation of a subset of CpG islands on the Xi^12,22^. The absence of SMCHD1 is also associated with DNA hypomethylation at its autosomal targets^17,21,25,27^. While these epigenetic changes have helped to place Smchd1 somewhere upstream of DNA methylation in the ontogeny of gene silencing, they have not revealed how Smchd1 functions to enable transcriptional repression.

At all of Smchd1’s genomic targets, chromosome conformation at different scales is known to play a role in regulating gene expression, exemplified by the Xi and the clustered protocadherins. On the Xi, two mega-domains much larger in size than TADs have been reported, which hinge on the clustered macrosatellite repeat *DXZ4*. In contrast, the active X (Xa) has a TAD structure similar to what is observed on autosomes^6,28−^^30^. Of note, the mega-domain structure does not appear to be critical for gene silencing, as deletion of *DXZ4* causes its collapse with no accompanying change in silencing of the Xi^28,30^. At the clustered protocadherins, short range enhancer-promoter chromatin looping occurs to regulate gene activation^31^. We previously reported that Smchd1 represses the protocadherins by binding to promoters and the enhancer at the cluster. We proposed a model based on the clustered protocadherin locus, where Smchd1 mediates chromosome conformation^17^, however we did not directly analyse conformation, nor did we consider the Xi because we used male cells. In addition to our own model, Nozawa *et al*. have also put forward a model for SMCHD1 regulating the chromosome structure of the human Xi. They found that the Xi in human 293 cells decompacted upon knockdown of SMCHD1, implicating SMCHD1 in the maintenance of a condensed Xi chromosome in human cells^26^.

Here we tested whether Smchd1 has a role in maintaining chromosome structure using *in situ* chromosome conformation capture followed by sequencing (*in situ* Hi-C). We report genome-wide changes to long-range chromatin interactions upon deletion of *Smchd1* in female cells, predominantly at Smchd1 bound loci. Disruption of long-range chromatin interactions at the *HoxB* locus prompted us to investigate the role of Smchd1 in regulating the *Hox* clusters, thereby implicating Smchd1 in the developmental regulation of *Hox* gene silencing. Remarkably, the widespread increase in X-linked chromatin interactions in the absence of Smchd1 did not result in the reactivation of genes from the Xi. Rather, we see an increase in intensity and occupancy of H3K27me3 on the Xi that is not associated with a loss of DNA methylation, suggesting that H3K27me3 enrichment is not due to enhanced recruitment of PRC2 to unmethylated CpG-rich regions^32-34^. These data suggest that Smchd1 may play a role in limiting the access of chromatin modifying proteins. Furthermore, genome-wide profiling of chromatin modifications and DNA accessibility suggests that Smchd1 may play a role in limiting long-range chromatin interactions that are permissive for transcription.

## Results

### Smchd1 is required for the maintenance of higher-order chromatin organisation at target loci

To test whether loss of Smchd1 alters long-range chromatin interactions at its targets, we performed *in situ* Hi-C in primary neural stem cells (NSCs) derived from the forebrains of E14.5 female *Smchd1*^fl/fl^ embryos, and in paired cells one week post Cre-mediated deletion of *Smchd1* (Fig. 1a). Conditional deletion of *Smchd1*^35^ in female NSCs allowed us to circumvent the female-specific embryonic lethality owing to a failure of X inactivation during development^12^ and to ask whether Smchd1 is involved in maintaining X-linked chromatin interactions in female cells. The three biological replicates for each genotype were highly correlated, and in multidimensional scaling analysis clustered by genotype, suggesting that higher order chromatin conformation is altered upon deletion of *Smchd1* (Supp. Fig. 1).

**Figure 1.**
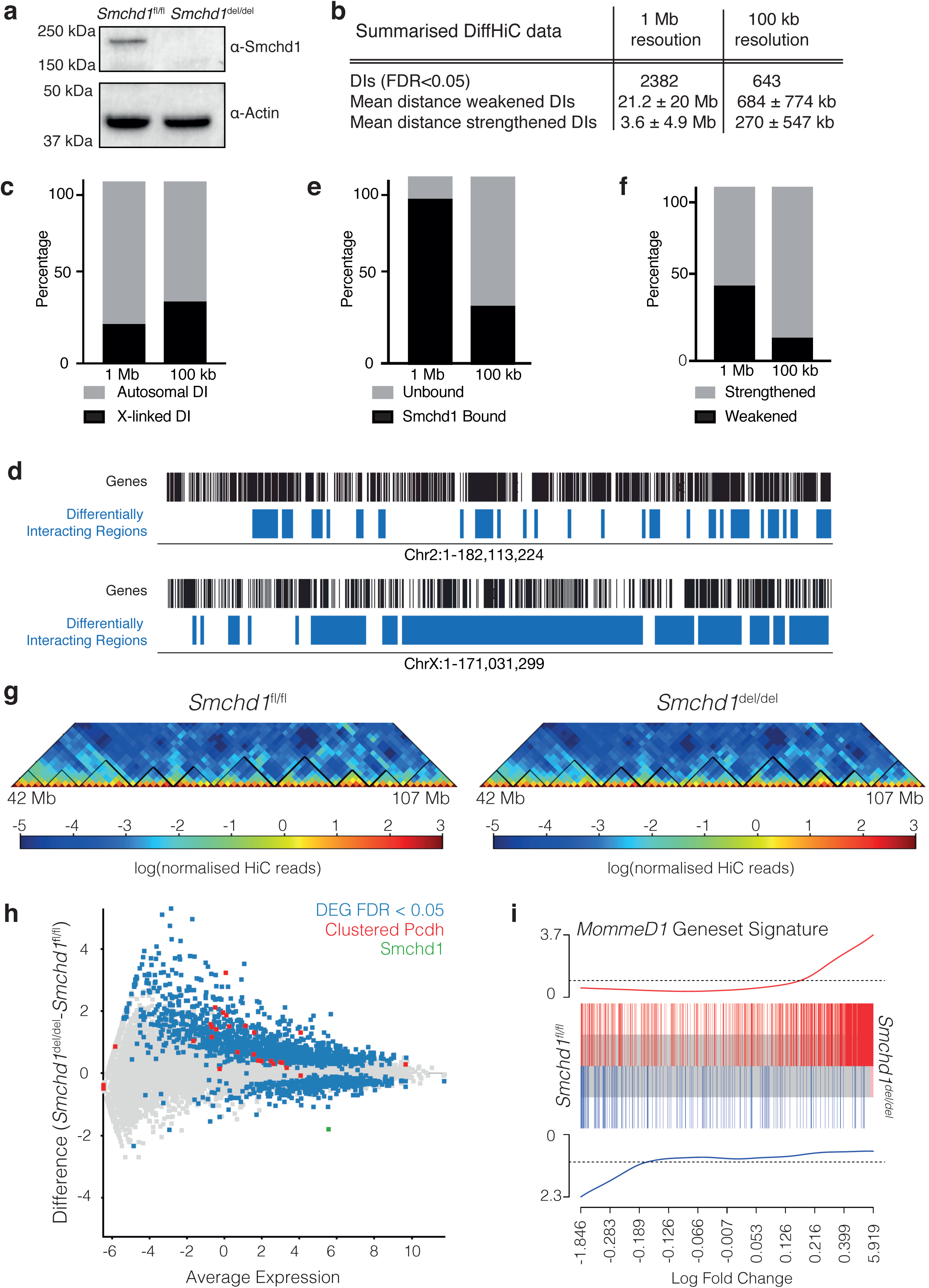
Smchd1 is required to maintain higher-order chromatin organisation at target loci, and to maintain gene silencing at a subset of its autosomal targets. **a.** Validation of Smchd1 deletion one week post Cre-mediated deletion in NSCs via Western blot for Smchd1, with Actin used as a loading control. **b.** Summarised data (mean ± standard deviation) from diffHic analyses between *Smchd1*^del/del^ and *Smchd1*^fl/fl^ NSCs at 1 Mb and 100 kb resolutions. n =3, FDR < 0.05. **c.** Bar graph showing the proportion of autosomal (grey) and X-linked (black) differential interactions. **d.** Distribution of individual differential interactions along the length of the X chromosome (bottom) with chromosome 2 shown above as a representative autosome for comparison. Genes are represented as black lines, and each 1 Mb DI anchor point is represented as a blue box. **e.** Bar graph showing the proportion of DIs that overlap a Smchd1 bound region of the genome (black) and those that do not (grey). **f.** Bar graph showing the proportion of DIs that are weakened (black) vs strengthened (grey) in the absence of Smchd1. Strengthened interactions refer to those that are detected more frequently in *Smchd1*^del/del^ samples than in *Smchd1*^fl/fl^ samples, whereas weakened interactions are those that are detected more frequently in *Smchd1*^fl/fl^ samples. **g**. Normalised Hi-C interaction frequencies at 100 kb resolution displayed as a heat map and rotated 45° for the 42 Mb to 107 Mb region on chromosome 11. Left panel is the *Smchd1*^fl/fl^ heatmap, right panel is *Smchd1*^del/del^ heatmap. The colour of the contact map, from blue to red, indicates the log_2_(contact frequency). Black triangles bound TADs detected by TADbit. **h.** Gene expression data from female *Smchd1*^fl/fl^ (n=3) (x axis) against *Smchd1*^del/del^ (n=3) samples (y axis). Genes that are differentially expressed between *Smchd1*^fl/fl^ and *Smchd1*^del/del^ samples (FDR<0.05) are shown in blue, clustered protocadherins in red and all other genes in grey. Axes are mean log_2_ counts per million (x) and log_2_ fold-change (y). **i.** Barcode plot of genes significantly upregulated (red bars, top of plot) and down-regulated (blue bars, bottom of plot) in *Smchd1*^MommeD1/MommeD1^ NSCs compared with wild-type, in the *Smchd1*^del/del^ versus *Smchd1*^fl/fl^ contrast. An enrichment line for each set shows the relative enrichment of the vertical bars in each part of the plot.

We used the bioinformatics package diffHic to ask whether there were differential chromatin interactions between *Smchd1*^fl/fl^ and *Smchd1*^del/del^ female NSCs. DiffHic uses edgeR statistics to perform a differential analysis of chimeric read counts that map to different genomic bins between experimental conditions^36^. To assess the extent to which loss of Smchd1 alters chromatin conformation, we performed analyses at two resolutions. EdgeR statistics are based on raw read counts and make statistical assumptions based on sequencing coverage, so we reasoned that analysis at 1Mb would afford us greater power to assess regions with fewer read counts; namely very long range interactions, as HiC interactions are known to decay as genomic distance increases^37,38^ (Supp. Fig. 1). diffHic analysis at 100kb resolution would then allow us to assess short range interactions between regions less than 1Mb apart that could not be measured in a lower resolution analysis (Supp. Fig. 2).

At 1 Mb resolution we detected 2382 significant differential interactions (DI) genome-wide (Fig. 1b). We were also able to detect these differential interactions at 500 kb resolution (87% overlap, Supp. Table 1). At increased resolution (100 kb), we detected 643 significant differential interactions between Smchd1 replete and deleted cells (Fig. 1b). We observed that many previously described Smchd1 target genes were found in the differential interactions from analyses at both resolutions, including the *Snrpn* imprinted cluster and three of the four *Hox* clusters (A, B and C), for which differential interactions at the *HoxB* cluster were most significant (Supp. Table 1). As expected given Smchd1’s role in XCI, X-linked differential interactions were over-represented (Fig. 1c, d).

We sought to determine whether changes to the chromatin that occur upon *Smchd1* deletion represent a disruption to the partitioning of the genome into A and B compartments, or the maintenance of TAD boundaries. Given Smchd1 occupancy is limited to a subset of autosomal binding sites, we did not expect to see global changes to the genome architecture, as when structural proteins such as Rad21 or Ctcf are depleted^8,9^. Analysis of the compartments revealed 90% of A and B compartments are maintained upon Smchd1 deletion (Supp. Fig. 2). Our HiC heatmaps indicate that TAD boundaries are unchanged in the absence of Smchd1 (Fig. 1g). Indeed, while genome-wide analysis of TAD boundaries detected changes in the strength of some TAD boundaries, the boundaries themselves are largely maintained (Supp. Table 2). These data suggest that Smchd1 does not play a role in the maintenance of compartments or TAD boundaries.

We analysed differential interactions in relation to Smchd1 occupancy in NSCs as determined by chromatin immunoprecipitation followed by high-throughput sequencing (ChIP-seq). We performed Smchd1 ChIP using our newly created *Smchd1*^GFP^ knock-in mice, which express a functional Smchd1-GFP fusion protein (Supp. Fig. 3). We generated high resolution ChIP-seq data under two distinct conditions: the first allowed for profiling of Smchd1 occupancy in accessible DNA by enzymatic fragmentation using MNase; the second profiled more inaccessible chromatin, including the Xi, using long sonication in non-ionic detergent (Supp. Fig. 3). Both methods resolved binding sites concordant with previously determined Smchd1 occupancy (Supp. Table 3)^17,26^. We then compared differential interactions with Smchd1 autosomal and Xi occupancy, and interestingly found that the two different resolutions potentially revealed something of biological interest. At 1Mb resolution we found 88.1% DIs are anchored in a region that contains a Smchd1 binding site, though this is only true for 31.4% DIs determined at 100 kb resolution (Fig. 1f). Intriguingly, while we detected long-range interactions that were both strengthened (43%) and weakened (57%) upon Smchd1 deletion at 1Mb resolution, at 100 kb resolution the vast majority of differential interactions are strengthened (Fig. 1g, Supp. Table 1). In agreement with these observations, analysis using the bioinformatics tool HiCCUPS^39^, which uses a local background model to identify short-range looping interactions at 10 kb resolution, also shows an increase in short-range interactions upon loss of Smchd1 (Supp. Table 4). To better understand the nature of these differential interactions, we sought to relate changes in higher order chromatin structure to transcription.

### Smchd1 is required for the maintenance of gene silencing at a subset of autosomal targets

Smchd1 has been shown to have an important role in gene silencing during development, but it is unknown whether Smchd1 is required for the maintenance of gene silencing. We therefore tested whether the widespread changes in genome organisation upon acute loss of Smchd1 were associated with changes in gene expression. RNA-seq performed on samples taken at the same stage as Hi-C revealed 463 significantly differentially-expressed genes (DEG) between *Smchd1*^fl/fl^ and *Smchd1*^del/del^ female NSCs (Fig. 1h, Supp. Table 5). Within this gene set were known Smchd1 targets, including clustered protocadherin genes and *Snrpn* cluster imprinted gene *Ndn*. We performed a gene set enrichment analysis using Roast and found that gene expression in *Smchd1*^del/del^ cells is concordant with male *Smchd1* null (*Smchd1*^MommeD1/MommeD1^) NSCs (p=0.0015, Fig. 1i). Analysis of gene expression data in relation to chromatin conformation found 69% of DEG fell within a differentially interacting region detected with diffHic (p = 0.00000164, hypergeometric probability). This list included the clustered protocadherin genes, which display both increased expression upon Smchd1 deletion, and also strengthened interactions determined by diffHic (Supp. Table 1), suggesting Smchd1 is required for the maintenance of chromosome architecture and gene silencing at some of its autosomal targets. By contrast, X-linked and *Hox* genes were not over-represented in the list of differentially-expressed genes (Supp. Table 5), indicating Smchd1 may not be required to maintain gene silencing in these cases. Taken together, our analyses have identified two classes of Smchd1 targets: those at which Smchd1 is important for maintaining both chromatin architecture and gene silencing; and those at which gene silencing is refractory to loss of Smchd1 in a differentiated cell, despite changes to higher order chromatin organisation.

### *Smchd1* deletion results in activation of autosomal enhancers

Our previous study found Smchd1 bound at distal *cis*-regulatory elements in male NSCs, but little is known about Smchd1 function in these regions. To better understand the effect of loss of Smchd1 on distal regulatory elements, we profiled DNA accessibility using Assay for Transposase Accessible Chromatin using sequencing (ATAC-seq). We identified 4205 differentially accessible peaks between *Smchd1*^fl/fl^ and *Smchd1*^del/del^ in male and female NSCs genome-wide (FDR<0.05, Supp. Table 6), where *Smchd1* deletion was associated with increased accessibility (Fig. 2a, b). We filtered the list of differentially accessible peaks for regions that differed in coverage by at least two-fold (913 peaks), to focus on regions at which we found the greatest change in accessibility upon loss of Smchd1.

**Figure 2.**
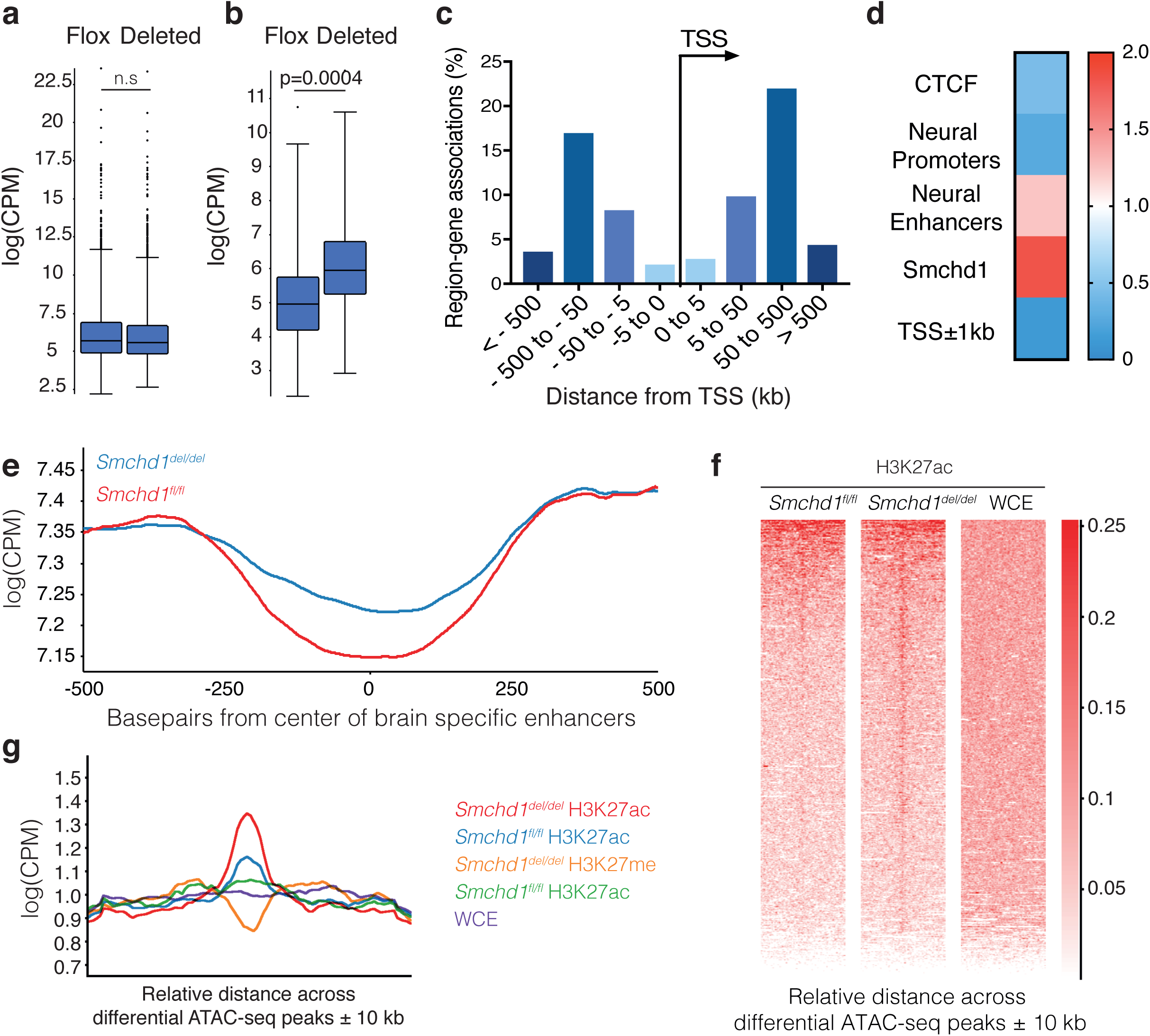
Smchd1 deletion results in activation of autosomal enhancers and increased promoter-enhancer interactions. **a-e**. Data from n=5, three female samples, and two males. **a/b.** Box plots showing coverage at all ATAC-seq peaks (**a**) and 4205 differential ATAC-seq (**b**) in log(cpm) in *Smchd1*^fl/fl^ and *Smchd1*^del/del^ NSCs. The box indicates the first to third quartile, horizontal line indicates the median, the whiskers indicate the median plus/minus the interquartile (25-75%) range multiplied by 2, and individual points that fall outside of this range are shown as circles. **c.** Distribution of the 913 differential ATAC-seq peaks that showed a greater than two-fold change in accessibility, relative to transcriptional start sites (TSS) calculated using the GREAT algorithm. The percentage of associations is plotted on the y-axis, and the distance from the TSS in windows as indicated in kb are plotted on the x-axis. **d.** Fold enrichment of chromatin features in 913 differential ATAC-seq peaks over the population. The colour scale indicates the fold enrichment. Smchd1 binding sites were defined in this study (Supp. Table 2) and all other genomic features were taken from Shen *et al.^40^***. e.** Coverage in log(cpm) of the 4205 differential ATAC-seq peaks centred over neural enhancers ± 500 bp. Reads from *Smchd1*^fl/fl^ in red and reads from *Smchd1*^del/del^ NSCs in blue. Note the truncated axis. **f.** Heat maps depicting H3K27ac signals and WCE control centred around the 913 differential ATAC-seq peaks ± 10,000 kb (x axis). y-axis presents the 913 differentially accessible peaks sorted by H3K27ac enrichment. n=1. **g**. Average H3K27me3 and H3K27ac over the 913 DAP shown in **e**, for *Smchd1*^fl/fl^ and *Smchd1*^del/del^ NSCs.

Whereas ATAC-seq reads tend to be highly enriched at the transcriptional start site (TSS) of genes, using the Genomic Regions Enrichment of Annotations Tool (GREAT), we found that the majority of the differentially accessible peaks sit 50-500 kb from the TSS (Fig. 2c). Consistent with this, we found differentially accessible peaks were enriched for E14.5 embryonic brain enhancers when compared with the population (132/913, Hypergeometric probability = 0.0047), but not promoters (33/913, Hypergeometric probability = 1, Fig. 2d)^40^. Looking at the distribution of differential reads over E14.5 embryonic brain enhancers, we observed a subtle increase in accessibility centred around these regions upon *Smchd1* deletion (Fig. 2e).

It is known that enhancer usage is highly tissue and cell-type specific^40^. We wanted to determine whether the remaining differentially accessible peaks might also contain non-neural enhancers that are becoming ectopically activated in the absence of Smchd1. Therefore, we performed ChIP-seq in *Smchd1*^del/del^ and control NSCs for H3K27ac, a mark found at active enhancers, and compared to H3K27me3, which is synonymous with inactive or poised enhancers. We analysed the regions under the 913 differentially accessible peaks and observed an increase in coverage of H3K27ac upon Smchd1 deletion (Fig. 2f, Supp. Table 7). The differentially accessible peaks also had a tendency to lose coverage of H3K27me3 upon loss of Smchd1 (Fig. 2g, Supp. Table 7). On occasion, the short-range interactions brought differentially expressed genes into proximity with H3K27ac marked regions. Taken together, these data suggest there could be ectopic activation of enhancers upon deletion of Smchd1.

### Smchd1 is involved in regulating *Hox* cluster conformation, and *Hox* gene silencing in development and differentiation

The most significant differential interaction from in situ Hi-C analysis at 1 Mb resolution was a lost interaction anchored at Chr11:96-97 Mb. This region contains the *HoxB* cluster, at which Smchd1 is highly enriched by ChIP-seq (Fig. 3a). In total we detected 23 weakened long-range interactions from the region containing the *HoxB* cluster upon deletion of *Smchd1*. Of note, 14/23 differential interactions were anchored in other clustered gene families that are Smchd1 targets, including the olfactory receptors and *Slc36* gene family (Fig. 3b, Supp. Fig. 4). Long range interactions were lost from the *HoxB* family in a range of 2–50 Mb. These differential interactions spanned a surprisingly large distance, suggesting that many are extra-TAD interactions, although we found the TAD boundaries remained unchanged (Fig. 3c, Supp. Table 2). The differential interaction with the lowest false discovery rate (FDR) was a weakening of an interaction between the *HoxB* containing region and the *Keratin* gene cluster, which is also occupied by Smchd1 (Fig. 3d, Supp. Fig. 4). We have validated this finding by DNA FISH, observing a decreased frequency of interaction between the *HoxB* and *Keratin* gene clusters post *Smchd1* deletion in NSCs (Fig. 3e, f, Chi-squared test p<0.0005). These data suggest Smchd1 mediates these long-range, extra-TAD interactions. Intriguingly, we also detect enhanced short range interactions with diffHic and the formation of a loop between the distal end of the *HoxB* cluster and the neighbouring region (Supp. Table 4); however these short-range interactions do not bring the *HoxB* cluster into contact with regions marked by H3K27ac.

**Figure 3.**
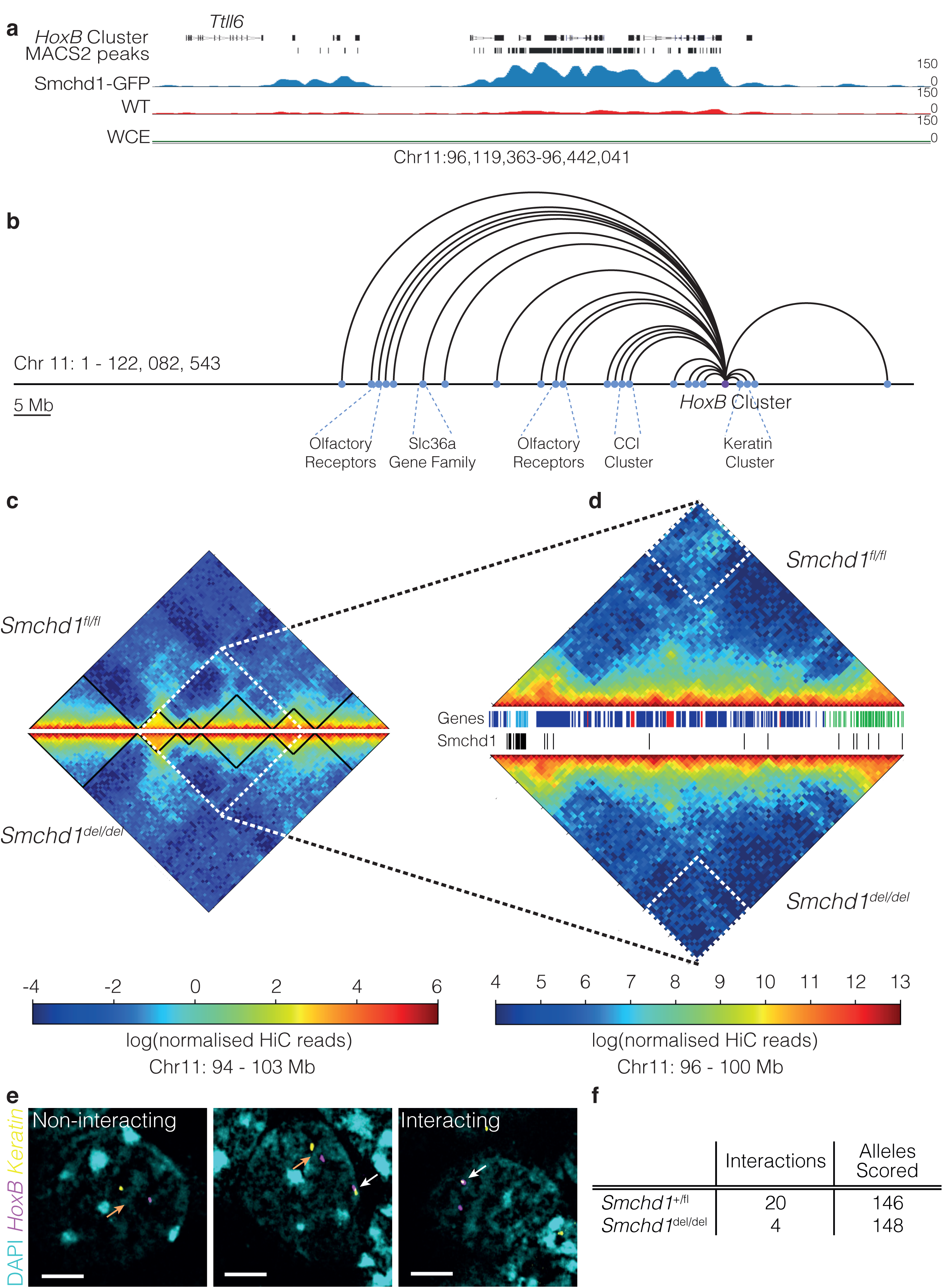
Smchd1 is required to maintain long-range interactions between the *HoxB* cluster and linked clustered gene families. **a.** ChIP-seq for Smchd1-GFP over the *HoxB* cluster. Genes are shown, followed by MACS2-like Smchd1-GFP ChIP-seq peaks as black bars. Smoothed Smchd1-GFP ChIP-seq track, normalised to reads from the whole cell extract, displaying positive enrichment of GFP ChIP-seq in *Smchd1*^GFP/GFP^ NSCs, alongside wild-type cells as an isotype control. Scale is normalised ChIP-seq reads. **b.** Schematic representation of *cis* differential interactions detected between female *Smchd1*^fl/fl^ and *Smchd1*^del/del^ NSCs at 1 Mb resolution at the *HoxB* cluster on chromosome 11. *HoxB* cluster is indicated in purple. Differentially interacting regions are indicated in blue. All interactions depicted are weakened in the absence of Smchd1. Half circles represent the interaction. **c.** Normalised Hi-C interaction frequencies at 100 kb resolution displayed as a heat map and rotated 45° for the region surrounding the differential interaction with the lowest FDR from diffHic analysis at 1Mb resolution, anchored at the *HoxB* and *Keratin* gene clusters. Above the horizontal is the *Smchd1*^fl/fl^ heatmap, below the horizontal *Smchd1*^del/del^. The colour of the contact map, from blue to red, indicates the log_2_(contact frequency). Black triangles bound TADs called by TADbit, dotted white region indicates the portion that is shown in higher magnification in **d.** Region of differential interaction in (**d.)** is depicted in a white dotted square. Between the heatmaps, genes are shown as bars (black, green for *Keratin* cluster genes, light blue for *HoxB* genes, red are differentially expressed in NSCs), Smchd1 ChIP-seq peaks are shown as black bars. **e.** DNA FISH using BAC probes against the *HoxB* cluster (magenta) and the *Keratin* gene cluster (yellow). DNA is stained with DAPI (cyan). The scale bar represents 4 *μ*m. Images show loci that are scored as not interacting (orange arrows, left and middle panels) or interacting (white arrows, middle and right panels). **f.** Quantification of number of *HoxB*-*Keratin* interactions from n=146 alleles n=148 alleles from *Smchd1*^fl/+^ and *Smchd1*^del/del^ cells, respectively.

Although we observed changes in both short and long-range chromatin interactions at the *Hox* gene clusters, and find Smchd1 highly enriched at all four *Hox* clusters in NSCs, we do not observe the upregulation of *Hox* genes upon *Smchd1* deletion. NSCs are derived from the embryonic forebrain, where *Hox* genes are not activated during development. Furthermore, epigenetic control of the *Hox* loci is regulated by multiple, redundant mechanisms in committed cell lineages, meaning NSCs may be an intractable system for studying perturbations to the maintenance of *Hox* gene silencing. To test if Smchd1-mediated higher order chromatin structures play a role in establishment or maintenance of developmental *Hox* gene silencing, we analysed the effect of *Smchd1* deletion in murine embryonic stem cells (mESC) differentiating into neuromesodermal progenitors (NMPs). mESC do not express *Hox* genes, but as they are directed to differentiate into neuromesodermal progenitors they turn on and off *Hox* gene expression in a way that mimics the collinear expression pattern of the tailbud in the developing embryo^41,42^.

One week post-Cre-mediated recombination, paired male *Smchd1*^fl/fl^ and *Smchd1*^del/del^ mESCs were differentiated using neuromesodermal progenitor protocols over eight days to activate the full complement of *Hox* genes^42,43^. Cells were sampled for RNA-seq at days 1, 5, and 8 of differentiation: timepoints selected to capture different stages of *Hox* gene activation and silencing (Fig. 4a). At days 1 and 5 there were 13 and 1 differentially-expressed genes between *Smchd1* deleted and replete samples, respectively, with Smchd1 itself the most significant differentially-expressed gene (Supp. Fig. 5). At day 8, however, there were 1425 differentially-expressed genes (Supp. Fig. 5, Supp. Table 8), which included known Smchd1 targets such as the clustered protocadherin genes and genes from the *Snrpn* imprinted cluster. Control samples behaved as expected, with anterior *Hox* genes activated by day 5 and posterior *Hox* genes at day 8. Interestingly, at day 8 in *Smchd1*^del/del^ neuromesodermal progenitors, 19 *Hox* genes were upregulated compared to controls (FDR<0.05). This included *Hox* genes from along the length of all four paralogous clusters (Fig. 4b). These data provide the first evidence that Smchd1 is required for *Hox* gene silencing during differentiation *in vitro*.

**Figure 4.**
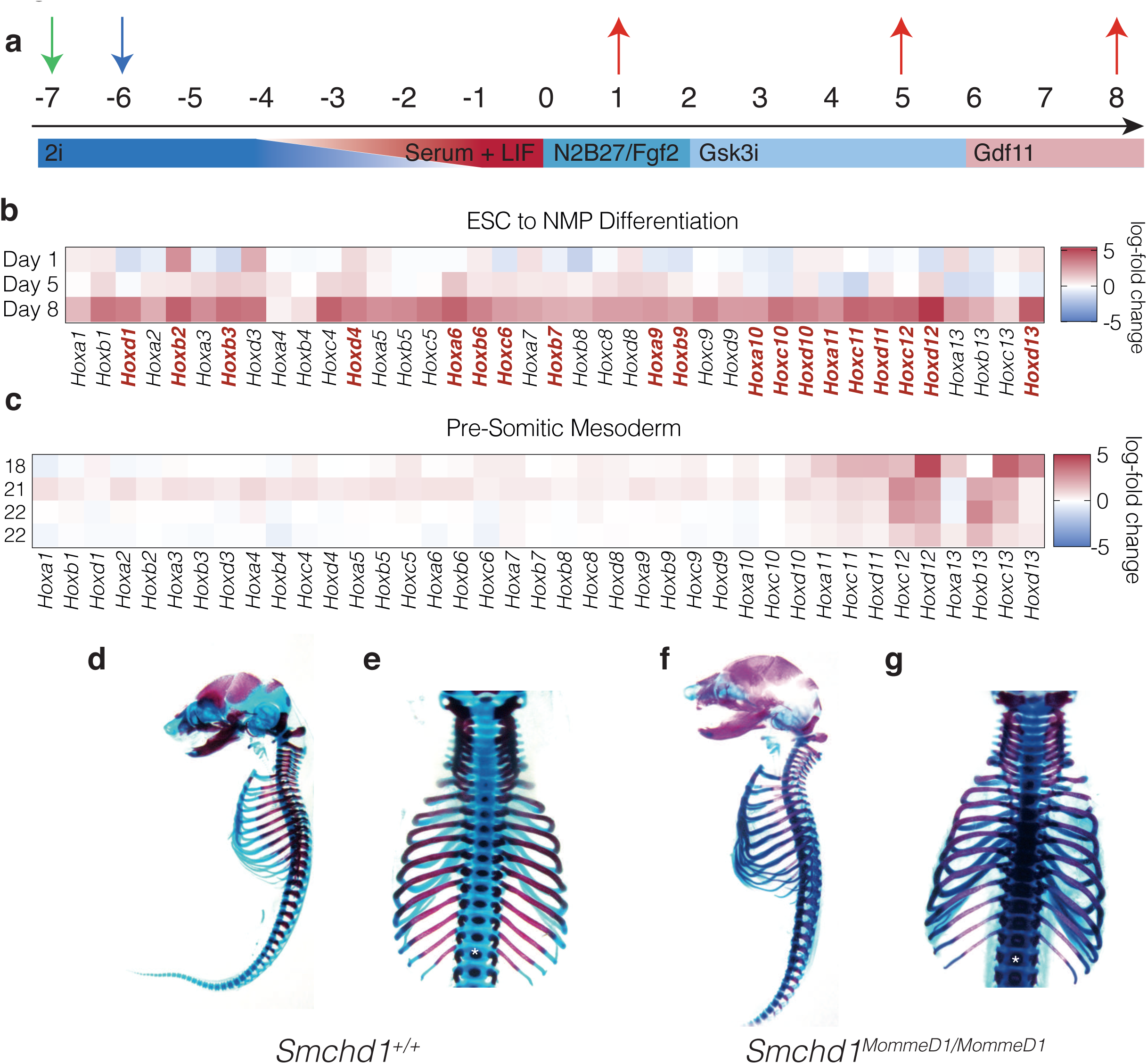
Smchd1 is required for developmentally regulated *Hox* gene silencing *in vitro* and *in vivo*. **a.** Schematic of *in vitro* differentiation of mESC to neuromesodermal progenitors. mESC are derived in 2i + LIF medium. One week prior to differentiation (D-7) cells were infected with MSCV-Cre-puro or MSCV-puro retrovirus (green arrow), and transduced cells selected with puromycin (blue arrow). Cells were weaned into 15% Serum + LIF medium. For differentiation cells were changed into N2B27 medium containing Fgf-2. At day 2 (D2) Gsk3i, and at D6 Gdf11 were added. Cells were harvested for RNA-seq at day (D) 1, 5 and 8 (red arrows). **b.** Heatmaps showing the log_2_(fold change) expression differences in *Hox* genes between *Smchd1* deleted and control samples at day 1, 5 and 8 of differentiation from mESC to neuromesodermal progenitors. Red to blue colour indicates fold change. Genes written in red have statistically significant differences at day 8, n=2. **c.** As for **b**, except samples are presomitic mesoderm from somite matched *Smchd1*^+/+^ and *Smchd1*^MommeD1/MommeD1^ E9.5 embryos. Somite numbers are indicated at the left. Log_2_fold change is shown for n=4 female *Smchd1*^MommeD1/MommeD1^ and n=3 female *Smchd1*^+/+^ embryos. **d/f.** Representative sagittal views of whole-mount E17.5 male *Smchd1*^+/+^ (**d**) and *Smchd1*^MommeD1/MommeD1^ (**f**) embryos with forelimbs and hindlimbs removed, stained with Alizarin red and Alcian blue for bone and cartilage, respectively. **e/g.** As per **d/f** showing dorsal view. Putative T13 is indicated with an asterix, note absence of rib on T13 in the *Smchd1*^MommeD1/MommeD1^ embryo, representing homeotic transformation.

As *in vitro* differentiation assays may not be subject to the same selective pressures and strict regulatory pathways employed during development, we wanted to test whether Smchd1 was involved in *Hox* gene silencing to the same extent *in vivo*. Previous work from our group and others identified *Hoxb7* and *Hoxb1* as differentially-expressed genes in microarray data for whole-embryos lacking Smchd1^23,25^; however, this approach was not designed to detect *Hox* gene changes and was limited based on the whole embryo samples. The presomitic mesoderm is a region in the embryo that continually expands and segments during development, giving rise to vertebral precursors known as embryonic somites. Patterning within the presomitic mesoderm is critically regulated by the collineaer activation of an anterior-to-posterior *Hox* code as development proceeds. We reasoned this would be a sensitive tissue to investigate the role of Smchd1 in *Hox* gene silencing, as has been performed for other *Hox* regulators^44^. We performed RNA-seq on the presomitic mesoderm of somite matched E9.5 *Smchd1*^+/+^ and *Smchd1*^MommeD1/MommeD1^ embryos (Supp. Fig. 6). We found premature activation of posterior *Hox* genes, including the *Hox 11, 12* and *13* paralogous cluster genes, but no differences in anterior *Hox* genes (Fig. 4c, Supp. Table 9). The discrepancy in anterior *Hox* regulation may stem from inherent difference between *in vitro* and *in vivo* conditions, or may reflect differences in the developmental state of Day 8 *in vitro* versus E9.5 *in vivo* samples. Nonetheless, both studies support a functional role for Smchd1-mediated chromosome conformation playing a role in *Hox* gene regulation.

To test the functional consequence of the *Hox* gene changes we observed in the absence of Smchd1, we performed whole-mount skeletal preparations from E17.5 *Smchd1*^MommeD1/MommeD1^ and *Smchd1*^+/+^ male embryos, as the effects of *Hox* gene dysregulation in mouse are most pronounced in the embryonic skeleton^45^. As expected, all wild-type embryos displayed 26 presacral vertebral elements, with no deviation from the normal configuration of C7-T13-L6 (10/10). In contrast, all *Smchd1* null embryos analysed exhibited a reduction in the number of presacral elements to 25, with varying changes in vertebral morphology centred around the thoraco-lumbar junction (10/10). In half of the cases (5/10) we observed a complete loss of the thirteenth thoracic element, with no concomitant gain of lumbar or sacral elements (C7-T12-L6). A variation of this phenotype was seen in 3/10 embryos, with a dramatically reduced rib process on T13 and loss of one lumbar element (C7-T13[reduced]-L5, Fig. 4d-g). The final two *Smchd1*^MommeD1/MommeD1^ embryos exhibited a similar partial rib reduction or complete rib loss of at T13, but with additional asymmetric changes at the lumbo-sacral junction. Three of the *Smchd1* null embryos additionally showed presence of an ectopic rib on one side of C7 (Supp. Fig. 6). Together, these data demonstrate an essential role for Smchd1 in formation of the vertebral column, with changes at the lumbo-sacral junction being viewed as either a loss of one element, or alternatively, serial homeotic transformation from the 20^th^ vertebral element (T13) onwards. Such posteriorizing transformations are consistent with precocious or enhanced posterior *Hox* activation^46,47^. Moreover, they are commonly seen in mice deficient for one or more polycomb group protein^48-51^, demonstrating a role for Smchd1 in *Hox* gene silencing *in vivo*.

Taken together, our data suggest that *in vivo* Smchd1 is involved in silencing posterior *Hox* gene expression prior to activation. Given we see perturbations to long range chromatin interactions at the *Hox* loci in *Smchd1*^del/del^ NSCs, we propose that developmental Smchd1-mediated *Hox* gene silencing likely occurs through regulation of the higher order chromatin structure.

### Acute loss of Smchd1 alters X chromosome conformation but does not result in reactivation of the inactive X chromosome

Much like *Hox* gene regulation, X inactivation is under strict developmental control, and once in the maintenance stage of silencing, reactivation requires disruption of multiple redundant epigenetic pathways. While we saw major changes in X-linked chromatin interactions following Smchd1 deletion, our RNA-seq data suggested that acute loss of Smchd1 did not result in a gross reactivation of the Xi in female NSCs. To ensure that we were not missing stochastic X reactivation, we also generated RNA-seq data from NSCs derived from female X_129_^XistΔA^X_Castaneus_; *Smchd1*^del/fl^ embryos, in which the *Xist*^Δ*A*^ allele ensures that the Castaneus X chromosome is the obligate Xi^52^. One week following Cre-mediated deletion of *Smchd1*, we profiled transcription by RNA-seq, and looked for expression of X-linked genes that mapped to the Castaneus X, indicative of X chromosome reactivation in the absence of Smchd1. We did not find any differentially expressed X-linked genes on the Xi or indeed on the Xa (Fig. 5a, b, Supp. Table 10). These data show that Smchd1-mediated higher order chromatin organisation is dispensable for the short term maintenance of X inactivation in female NSCs.

**Figure 5.**
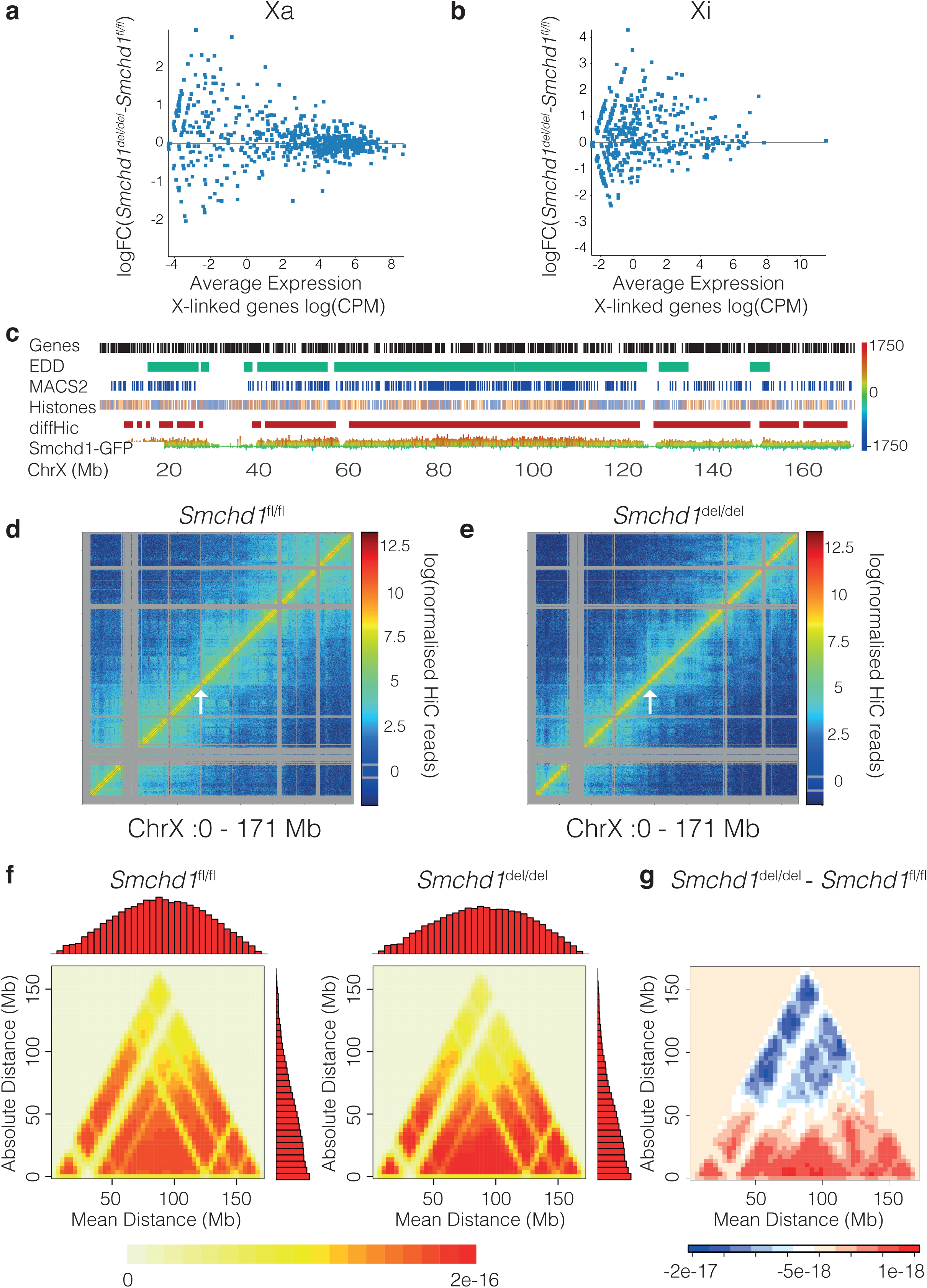
Smchd1 deletion appears to alter the conformation of the Xi, but is not required to maintain gene silencing on the Xi. **a-b.** X-linked gene expression data from female *Smchd1*^del/fl^ (n=2) (x axis) versus *Smchd1*^del/del^ (n=2) samples (y-axis). X-linked genes are represented in blue; no differentially expressed genes were detected on the Xi or Xa between *Smchd1*^del/fl^ and *Smchd1*^del/del^ samples (FDR<0.05) **a.** Plot for the 129-derived *Xist*^ΔA^ allele containing Xa **b.** Plot for the Castaneus-derived obligate Xi. **c**. Enrichment of Smchd1-GFP over whole cell extract on the X Chromosome. Height and colour of peaks indicates strength of read counts normalised to whole cell extract; red is most enriched, blue is most depleted. Position of MACS2-like peaks called using Seqmonk are indicated in blue. Enriched domain detector (EDD) domains are shown in green. n=2. Histones track depicts H3K27me3 (salmon) and H3K9me3 (blue) derived from Keniry *et al*.^54^. DiffHic track depicts differential interactions from our own analyses, n=3. **d-e.** Normalised Hi-C interaction frequencies at 100 kb resolution displayed as a heat map for the X chromosome in *Smchd1*^fl/fl^ (**c**) and *Smchd1*^del/del^ (**d**) NSCs. The colour of the contact map, from blue to red, indicates the log_2_(contact frequency). The arrow indicates *DXZ4*. **e.** A 45^°^ rotated transformed interaction matrix displayed as a heatmap for the X chromosome in **c-d** that shows the distribution of absolute distances against the mean distances of all interactions in the original matrix. The colour from yellow to red indicates the density estimated from the absolute and mean distances. **f.** Differential plot of the transformed interaction matrices for the X chromosome in *Smchd1*^del/del^ minus *Smchd1*^fl/fl^ in part **e**. The colour from blue to red indicates the difference between the two densities estimated in part **e**.

Previous work has suggested that the maintenance of the mega-domain structure of the Xi may not be required for gene silencing, because deletion of the *DXZ4* macro-repeat array dramatically altered the Xi architecture, but did not result in failed XCI during differentiation or X chromosome reactivation in differentiated cells^28,30^. We therefore used our Hi-C data to generate interaction maps for the X chromosome in *Smchd1* deleted and replete samples^53^. While we do not have the polymorphic samples required to detect genetic differences between the Xa and Xi in the Hi-C datasets, Smchd1 is not found on the active X chromosome in males^17^, and our Smchd1-GFP ChIP-seq reveals Smchd1 covering the majority of the presumptive Xi (Fig. 5c), similar to the report of SMCHD1 ChIP-seq in human cells^26^. Smchd1-GFP is notably absent from the gene-poor region that shows highest enrichment for H3K9 methylation^54^: a region where we observe no differential interactions. Based on Smchd1’s strong enrichment on the Xi, we can assume changes in X-linked chromosome architecture post-*Smchd1* deletion relate to the Xi. We also performed *in situ* HiC followed by analysis with diffHic in paired male samples, and found the overrepresentation of X-linked DIs to be specific to female samples, further suggesting that X-linked changes detected in our analyses reflect changes to the architecture of the Xi (Supp. Table 1). In the control samples, the interaction map is consistent with the known mega-domain structure of the Xi, hinged at *DXZ4* (Fig. 5d)^6,28−^^30^. In the absence of Smchd1, the mega-domain structure is less apparent (Fig. 5e), which is also observed in heatmaps from subsampled data (Supp. Fig. 7). However, analysis with diffHic showed *Smchd1* deletion results in strengthening of short-range interactions concomitant with loss of the long-range mega-domain structure (Fig. 5f, g). These data are consistent with a role for Smchd1 in limiting short range interactions on the Xi, and contributing to the unique structure of the Xi.

### Smchd1 is not required to maintain Xi chromosome compaction or accessibility in mouse

Given the role of SMCHD1 in the compaction of the human Xi^26^, we wanted to determine whether the observed large-scale changes in X-linked chromatin interactions upon *Smchd1* deletion were indicative of a loss of Xi compaction in mouse. We performed DNA FISH using an X chromosome paint in female *Smchd1*^fl/fl^ and *Smchd1*^del/del^ NSCs. The chromosome paint does not discern between Xa and Xi, but based on the reported 20-30% difference between the volume of the Xi and the Xa within individual nuclei^55^ we assumed the smaller of the two X chromosomes within each nucleus was the Xi. Our measurements were consistent with published data, indicating an approximate 30% difference between the volume of the two X chromosomes within individual nuclei in both *Smchd1*^fl/fl^ and *Smchd1*^del/del^ female NSCs. There was no difference in the mean volume of the presumptive Xi or Xa in female *Smchd1*^fl/fl^ and *Smchd1*^del/del^ NSCs in a population of cells, nor was there a change in the difference of the volumes of the two X chromosomes measured within individual nuclei (Fig. 6a-c, Supp. Fig. 7). Consistent with this, we did not see female-specific changes in X-linked chromatin accessibility, as determined by ATAC-seq in female *Smchd1*^fl/fl^ and *Smchd1*^del/del^ NSCs, when compared with males (Supp. Figure 7, Supp. Table 6). Together these data suggest that Smchd1 is not involved in maintaining chromatin compaction of the murine Xi.

**Figure 6.**
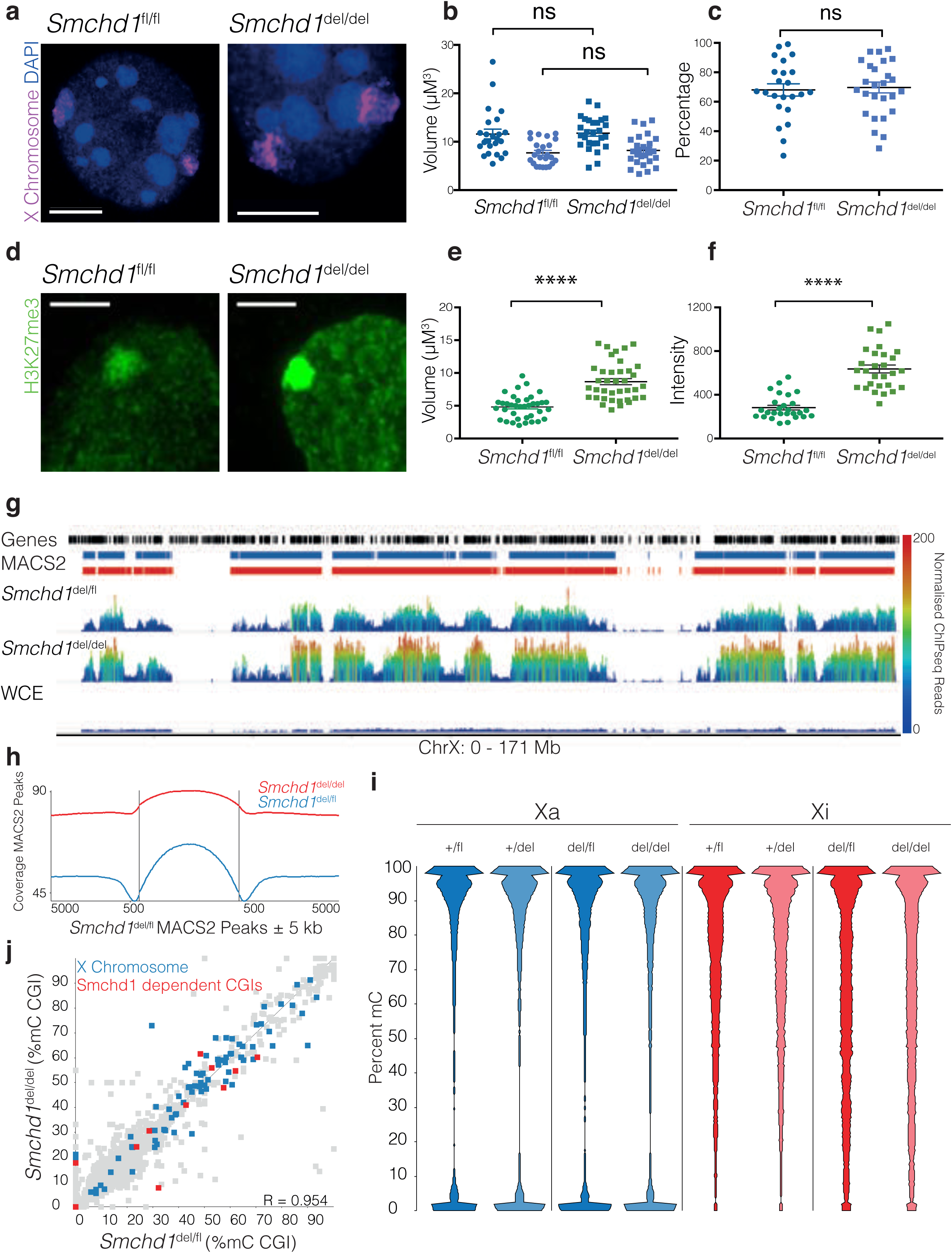
Smchd1 limits H3K27me3 spreading on the Xi, though it is not required to maintain Xi compaction. **a.** Confocal imaging with Airyscan processing of *Smchd1*^fl/fl^ (left) and *Smchd1*^del/del^ (right) NSC nuclei following DNA FISH with X chromosome paint, n=23 observations per genotype, n=2 cell lines. DAPI stained DNA is shown (blue) and X chromosome paint (magenta), shown with 3D rendering to measure volume, below confocal images. **b-c.** Dot plots showing the volumes of largest and smallest X chromosome in each nucleus (**b**) or percent difference between the volumes of largest and smallest X chromosome in each genotype (**c**). Statistical comparisons were made using a two-tailed student’s t-test, line indicates mean, error bars are SEM. **d.** Confocal images with Airyscan processing of H3K27me3 immunofluorescence in female *Smchd1*^fl/fl^ and *Smchd1*^del/del^ NSCs (green). **e.** Dot plots of volume and **f.** mean intensity of focal H3K27me3 enrichment in 3D using Imaris Statistical comparisons were made using a two-tailed student’s t-test, line indicates mean, error bars are SEM. Measurements from 38 observations in n=3 cell lines, p <0.00001. **g.** Genome browser tracks for the H3K27me3 ChIP-seq on the obligate Castaneus-derived Xi chromosome in *Smchd1*^del/fl^ and *Smchd1*^del/del^ NSCs (n=2). Genes are shown in black, MACS2-called peaks in *Smchd1*^del/fl^ NSCs are shown in blue, and MACS2-called peaks in *Smchd1*^del/del^ NSCs are shown in red. ChIP-seq tracks are shown for the active X chromosome (Xa) and inactive X chromosome (Xi) from *Smchd1*^del/fl^ and *Smchd1*^del/del^ NSCs (n=2). Scale is normalised ChIP-seq read counts, normalised for read depth using the total autosomal read count. **h.** Coverage of MACS2-called H3K27me3 peaks in both *Smchd1*^del/del^ and *Smchd1*^del/fl^ samples, centred around MACS2-called H3K27me3 peaks detected on the Xi of *Smchd1*^del/fl^ samples ± 5000 kb. **i.** Bean plots showing DNA methylation (%mC) on the Xa (blue) and Xi (red) as determined by RRBS in *Smchd1*^del/del^ (n=1), *Smchd1*^del/fl^ (n=1), *Smchd1*^del/+^ (n=1) and *Smchd1*^fl/+^ (n=1) NSCs. CpG analysed were those with a minimum coverage of 10 reads in each condition. The former 2 genotypes were derived from embryos haploinsufficient for Smchd1 during embryogenesis, while the latter 2 were Smchd1 replete. **j.** Dot plot showing the DNA methylation of CpG Islands (%5mC) that map to the castaneus genome in *Smchd1*^del/fl^ (x-axis) and *Smchd1*^del/del^ NSCs. CGIs on the Xi are indicated in blue. CGIs that are hypomethylated during development in female embryos in the absence of Smchd1 are indicated in red. Pearson’s correlation value (R) is shown on the bottom, right of the scatter plot.

### Loss of Smchd1 results in widespread H3K27me3 enrichment on the Xi

It was intriguing that extensive loss of long-range chromatin interactions on the Xi was not accompanied by concurrent alterations in gene expression, compaction, or chromatin accessibility. We were therefore interested in whether we could observe changes to an alternative silencing pathway on the Xi in the absence of Smchd1. We performed immunofluorescence for H3K27me3, the mark catalysed by PRC2, in female *Smchd1*^fl/fl^ and *Smchd1*^del/del^ NSCs (Fig. 6d). Surprisingly, upon Smchd1 deletion, there was a 2.25-fold increase in H3K27me3 intensity on the Xi and an increase in volume (Fig. 6d-f, Paired t-test, p<0.00001). We were also able to detect an enrichment of H3K27me3 on the Xi by ChIP-seq in female X_129_^XistΔA^X_Castaneus_; *Smchd1*^del/del^ NSCs compared with controls (Fig. 6g, Supp. Table 11). In addition to an increase in H3K27me3 signal on the Xi, *Smchd1* deletion also resulted in detection of H3K27me3 MACS2 peaks in regions where they would not normally be found on the Xi (Fig. 6g, h, Supp. Fig. 7), indicating spreading of H3K27me3. These data prompted us to re-examine H3K27me3 on the Xi in *Smchd1*^MommeD1/MommeD1^ female nuclei, and interestingly, there too we saw increased intensity of H3K27me3 staining on the Xi (Supp. Fig. 7)^12^, suggesting this change is not exclusive to acute loss of Smchd1. While the Xi enrichment of H3K27me3 is striking, we did not observe enrichment of this mark at Smchd1’s autosomal targets. Spreading of H3K27me3 domains on the Xi in the absence of Smchd1 has not been reported before.

One possible explanation for H3K27me3 enrichment in the absence of Smchd1, is that DNA hypomethylation could facilitate PRC2 recruitment^32-34^. Indeed, female embryos heterozygous or homozygous for a null allele of *Smchd1* display a dose-dependent reduction in DNA methylation at CGIs on the X chromosome^12,22^. Therefore, we assessed DNA methylation in NSCs derived from both X_129_^XistΔA^X_Castaneus_; *Smchd1*^+/fl^ and X_129_^XistΔA^X_Castaneus_; *Smchd1*^del/fl^ female embryos, one week post Cre-mediated deletion of *Smchd1*, using reduced representation bisulphite sequencing (RRBS). As expected^12^, female cells that have undergone XCI with a heterozygous deletion of *Smchd1* displayed reduced DNA methylation on the Xi (Fig. 6i, Supp. Table 12). However, we did not detect hypomethylation on the Xi upon acute loss of Smchd1, in cells either heterozygous or homozygous for *Smchd1* deletion, either at all CpGs or CpG islands (Fig. 6i, j, Supp. Table 12). Therefore an increase in H3K27me3 occurs independently of DNA hypomethylation following acute depletion of Smchd1. Taken together, our data raises the possibility that Smchd1 contributes to the maintenance of the local chromatin environment at target loci, potentially by blocking the access of other epigenetic regulators.

## Discussion

Smchd1 has well characterised roles in developmental gene silencing in both mouse and human^12,21-23,25,56,57^. In this study we have implicated Smchd1 in the developmental silencing of another fundamentally important set of genes, the *Hox* cluster genes. We present a genome-wide study of gene expression and chromatin architecture upon acute loss of Smchd1. Our data implicate Smchd1 in the maintenance of gene silencing and higher order chromatin organisation of target loci. Intriguingly loss of Smchd1 had a much more dramatic effect on the maintenance of chromatin conformation and composition than it did on accessibility and transcription. While *Smchd1* deletion resulted in a change in conformation and expression at the clustered protocadherins and imprinted targets, the same was not true for the inactive X chromosome. During development, Smchd1 is critical for X-linked gene silencing later in the ontogeny of X inactivation, which could implicate Smchd1 in the maintenance of X-linked gene silencing; however we have shown that dramatic remodelling of the X chromosome upon acute loss of Smchd1 is not associated with reactivation of genes from the silent X chromosome. We made similar observations at three paralogous *Hox* clusters. These findings show that select genes have a strictly developmental requirement for Smchd1 in regulating transcriptional silencing. Additionally we identify a subset of autosomal genes that require Smchd1 for the maintenance of gene silencing.

While we and others have previously suggested that Smchd1 may have a role in regulating higher order chromatin architecture, Smchd1’s role in chromatin structure had not specifically been explored^17,26,58^. By performing *in-situ* Hi-C, followed by powerful differential analyses with diffHic, we were able to determine that loss of Smchd1 does indeed result in genome-wide changes to long-range chromatin interactions. Analyses at two resolutions saw long-range interactions that largely reflect Smchd1 occupancy are weakened, whereas shorter-range interactions that are not enriched for Smchd1 occupancy are strengthened (Fig.7 a, b). Further examination of specific loci, and investigation of the changing chromatin landscape, along with gene expression, led us to propose that Smchd1 may function to maintain both chromatin architecture and gene expression, by limiting access of other epigenetic complexes that alter the chromatin. This is illustrated by our previous observation that the absence of Smchd1 at the clustered protocadherin locus results in an increase in Ctcf occupancy^17^. We now report that this too is true for H3K27me3 on the Xi, although at a much larger scale. Taken together, these data suggest that Smchd1 has the capacity to function at the chromatin as an insulating protein (Fig. 7b, c). It would be interesting to determine whether Smchd1 possesses insulating potential in other contexts, and towards other chromatin modifying proteins. The zinc finger protein Ying-Yang 1 (Yy1) has been shown to mediate developmentally regulated promoter-enhancer contacts in neural progenitor cells^59,60^. Potentially Smchd1 represses distal regulatory elements in NSCs by limiting Yy1-mediated promoter-enhancer contacts. Given the tissue-specific manner in which enhancers are regulated, it will be informative to study Smchd1 occupancy and its effect on the chromatin architecture in cells derived from different tissues, such as those where SMCHD1 plays an important role in disease^56-58,61,62^.

**Figure 7.**
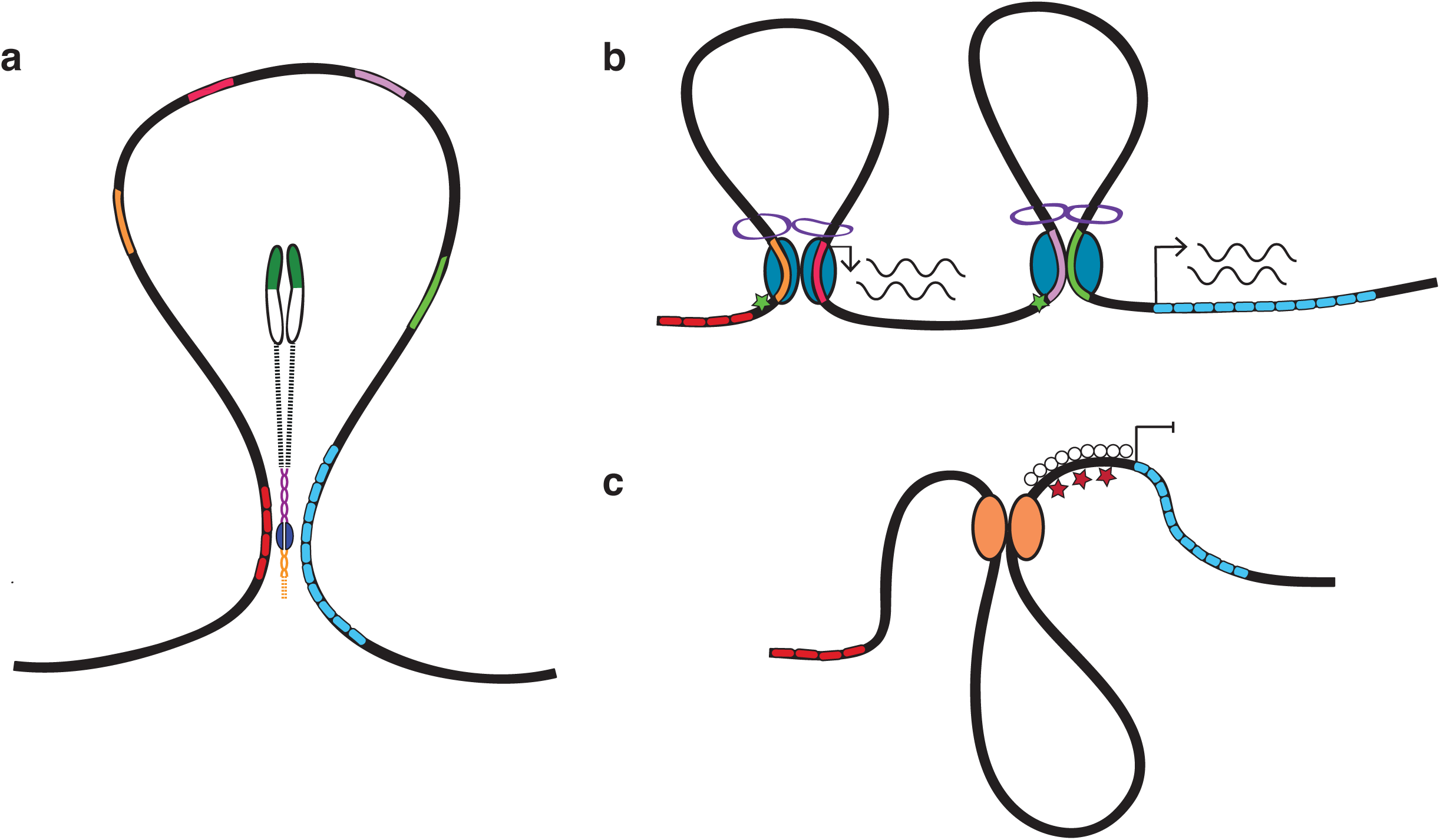
An insulating model for Smchd1 function: Smchd1 is involved in the maintenance of long-range repressive chromatin structures, which limit promoter-enhancer interactions that are permissive for transcription, and access to chromatin modifiers. **a.** In *Smchd1*^+/+^ cells, Smchd1 is targeted to regions of the genome that are repetitive in nature (red and blue repeating blocks), such as clustered gene families, including the *Hox* clusters, where it enables long-range chromatin looping. **b.** In the absence of Smchd1, long-range repressive interactions are lost, which based on our previous data allows for the formation of Ctcf-mediated (blue ovals) promoter-enhancer interactions that are permissive for transcription (depicted in purple-green and red-orange). **c.** On the inactive X, loss of Smchd1 results in enhanced H3K27me3. We propose this is due to loss of Smchd1’s capacity to insulate the chromatin from other chromatin modifiers (orange circles).

DiffHic analysis revealed a loss of long-range autosomal chromatin interactions upon Smchd1 deletion, suggesting that Smchd1 is involved in maintaining long-range chromatin interactions (Fig. 7a). This is illustrated by the *HoxB* locus, where interactions are weakened between the *HoxB* cluster, and a number of other clustered gene families occupied by Smchd1 on chromosome 11. These lost interactions occur across topologically associated domain (TAD) boundaries at a scale of tens of Mb. Moreover, we do not detect changes to TAD boundaries in these regions. Together, these data provide evidence that like a number of other epigenetic regulators, Smchd1 may be involved in mediating long-range chromatin interactions at target loci, potentially forming Smchd1 clusters such as those observed between H3K27me3 targets^63^. By focusing on the *HoxB* cluster, we found that in tissue restricted cells, there was a decoupling of a loss of Smchd1-mediated chromatin interactions, and transcriptional regulation at the locus; however by studying gene expression more dynamically *in vitro* and *in vivo*, we identified a role for Smchd1 in *Hox* gene silencing. Based on these data, we propose that Smchd1-mediated longrange interactions are repressive in nature, and function in developmentally regulated silencing pathways. Given the dynamic regulation of the chromatin architecture at the *Hox* clusters, it will be of interest to study Smchd1-mediated chromatin interactions in the context of collinear regulation of *Hox* gene expression during development, to better place Smchd1-mediated longrange interactions in the hierarchy of developmental gene silencing.

Given the prominent role of Smchd1 in XCI, it was no surprise to find X-linked regions were over-represented in the differentially interacting regions. While we could not discern between the Xi and Xa genetically in our in situ Hi-C data, the absence of Smchd1 from the male X chromosome, and the Xi enrichment of Smchd1 throughout the maintenance of XCI suggest that X-linked changes detected using diffHic arise from the Xi. The inactive X chromosome is unique in its higher order conformation; whereas the Xa is structured like an autosome, enriched for Ctcf and comprising many TADs, the Xi is devoid of local structures and rather is organised into two large mega-domains^28,30^. Our analyses suggest that upon Smchd1 deletion, the Xi adopts a structure that more closely resembles the structure of the Xa. Loss of Smchd1 and associated long-range chromatin interactions did not result in female-specific changes in chromatin accessibility, or the compaction of the murine X chromosomes as measured by DNA FISH. This is in contrast to observations by Nozawa *et al*., who suggested that SMCHD1 is involved in the compaction of the human Xi by bringing together H3K27me3 and H3K9me3 heterochromatin domains through an HBiX-HP1 pathway^26^. Our data suggest that in mouse, Smchd1 functions through a divergent pathway to regulate chromatin structure on the Xi. This is supported by Brideau *et al*., who found that the interaction between Smchd1 and the murine HBiX homologue Lrif1 is not conserved on the Xi in mouse^18^. An alternative possibility is that Smchd1 may play a role in maintaining the structure of the Xi, by insulating Ctcf-mediated structures that are associated with the Xa.

In conclusion, we have identified a role for the non-canonical SMC protein Smchd1 in the maintenance of higher order chromatin structures. We propose that Smchd1-mediated long range interactions have insulating capabilities, that limit access to other factors including chromatin modifying complexes and chromatin proteins that enable structures permissive for transcription. Given the important developmental role of SMCHD1 in mouse and human, it remains of interest to determine how Smchd1 is targeted to genes on autosomes and the inactive X chromosome to perform its functions as a structural protein.

## Methods

Methods and any associated references are available in online methods.

## Accession Codes

All genomics data has been deposited in the Gene Expression Omnibus, under accession number GSE111726.

## Acknowledgements

We thank Hannah Coughlin, Rhys Allan and Tim Johansson for useful discussions. This work was funded by the Australian National Health and Medical Research Council grant to MEB, JMM and MER (GNT1098290), and fellowships to JMM (GNT1105754) and MER (GNT1104924). NJ was supported by an Australian Research Training Program Fellowship. MEB was supported by a Bellberry-Viertel Senior Medical Research Fellowship. The Australian Regenerative Medicine Institute is supported by grants from the State Government of Victoria and the Australian Government. This work was made possible through Victorian State Government Operational Infrastructure Support and Australian National Health and Medical Research Council Research Institute Infrastructure Support Scheme.

## Author Contributions

NJ designed experiments, performed experiments, interpreted and analysed data and wrote the paper. AK designed experiments, performed experiments, interpreted and analysed data. MT, PH and MER contributed to bioinformatic analyses of the data. TB, KB and MI performed experiments. HB designed and performed experiments. EM designed experiments, performed experiments and interpreted and analysed data. GFK, IDT and AWM designed and performed experiments, JMM designed experiments, interpreted data and edited the paper. MEB designed experiments, interpreted data, wrote and edited the paper.

## Figure legends

**Supplementary Figure 1.**
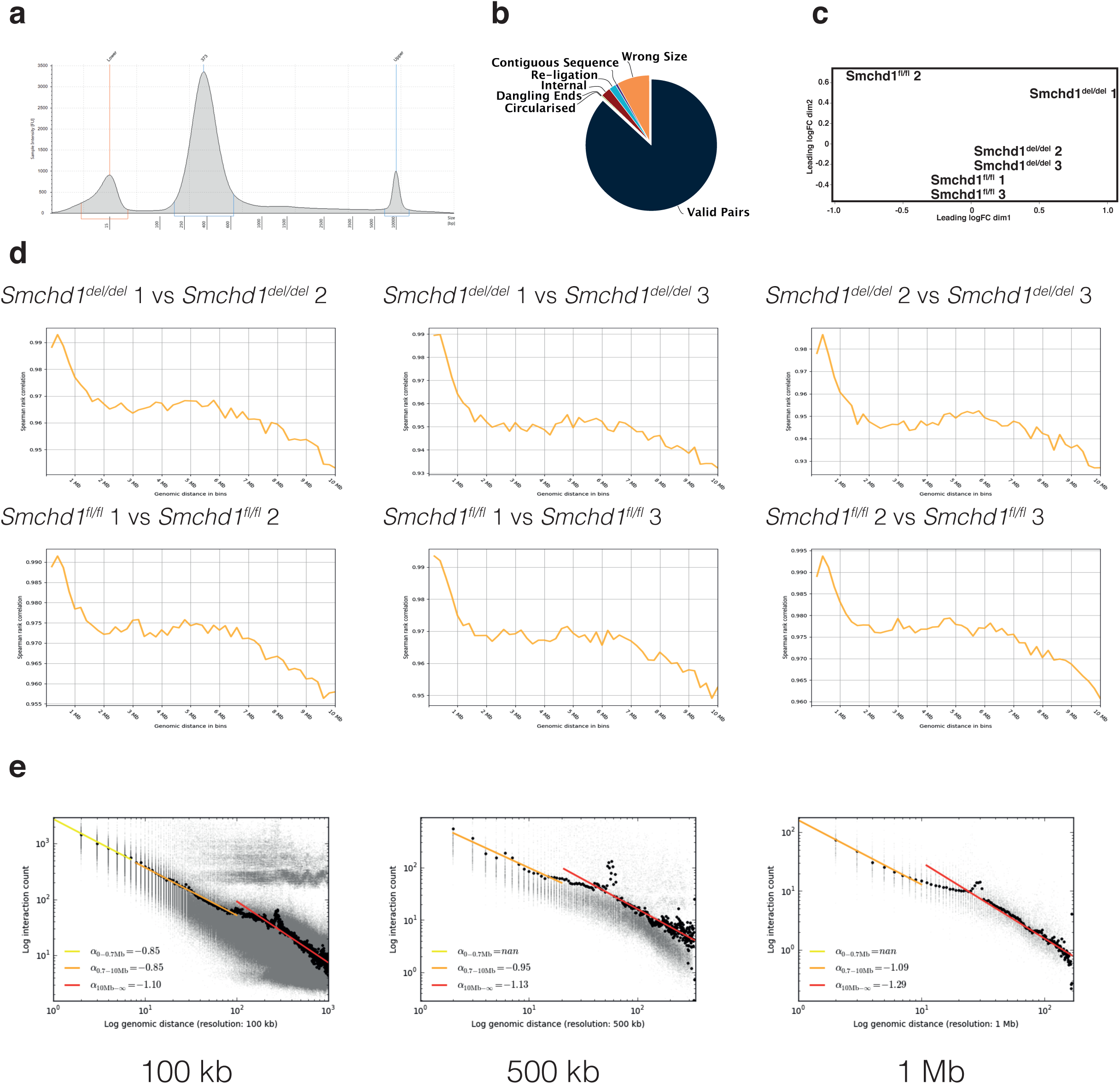
In situ Hi-C library quality control measures. In situ Hi-C was performed in n=3 *Smchd1*^fl/fl^ and *Smchd1*^del/del^ female NSC lines, one week post Cre-mediated deletion of *Smchd1*. **a.** Tapestation trace showing read-length distribution of pooled in situ HiC libraries run on a high-sensitivity D5000 Tape. Libraries were produced in the correct size range. **b.** These libraries were of good quality, as determined by HiCup. Pie graph from HiCup processing report representing the relative distribution of in situ HiC reads, following optimisation of restriction digest and conditions to prevent aggregation of nuclei. Valid pairs, or chimeric HiC reads, are represented in navy blue. All other colours represent a common experimental artefact. Notably the valid reads now make up 86% of the total reads. **c.** Multidimensional scaling (MDS) plot showing the relationship between the 3 *Smchd1*^fl/fl^ and 3 *Smchd1*^del/del^ in situ Hi-C samples. Samples separate by genotype on the first MDS component: Leading logFC dim1 (X axis); Leading logFC dim1: second MDS component (Y axis). **d.** High correlation between biological replicates indicated by a Spearman rank correlation between bins in two matrices at increasing genomic distances. **e.** The log interaction count (Y axis) compared to log genomic distance (X axis) shown for the 100 kb, 500 kb and 1 Mb resolution analyses. The trend line is shown for 3 different log genomic distances. These plots show the distance-dependent decay of HiC interaction frequency.

**Supplementary Figure 2.**
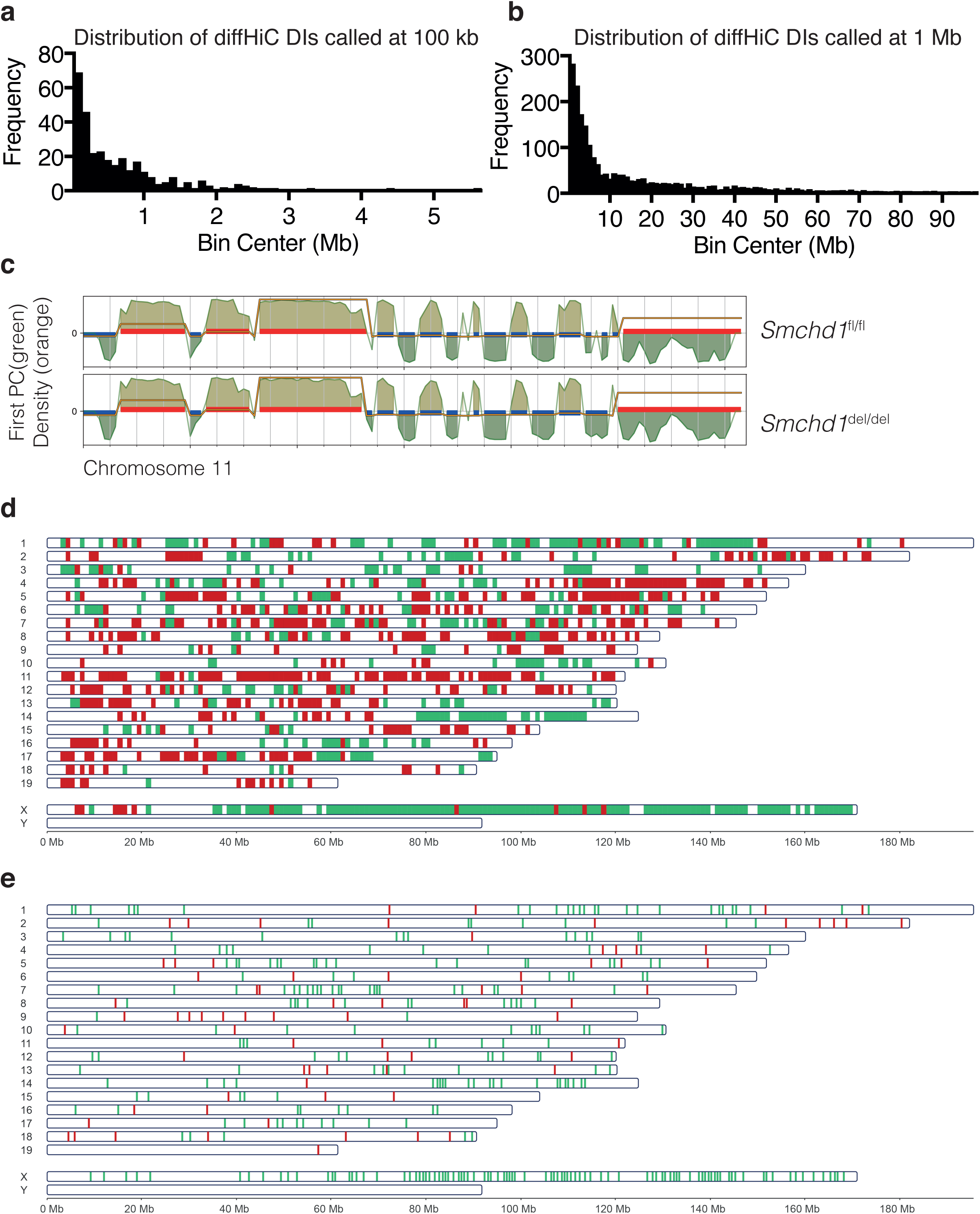
*In situ* HiC diffHiC differential interactions and compartment analysis. **a-b.** Histograms of the differential interactions (DIs) called at 100kb (**a**) and 1Mb (**b)**, plotting the distance between the two genomic anchors (Bin centre, X axis) against the frequency of interactions (Y axis). **c.** Summary TADbit analysis using the first eigenvector to identify compartments from a correlation matrix of normalised HiC data at 1MB resolution across chromosome 11 in *Smchd1*^fl/fl^ and *Smchd1*^del/del^ samples, showing no significant changes in compartment assignment (90% overlap genome-wide). The first principle component of the PCA on the Hi-C correlation matrix is shown as a green line, the density as an orange line, and the blue or red block indicates A vs B compartment **d-e.** Distribution of strengthened (green bars) and weakened (red bars) interactions at 1 Mb (**d**) and 100 kb (**e**) resolution. Differential interactions are spread across all chromosomes, but cluster at Smchd1 binding sites such as the X chromosome.

**Supplementary Figure 3.**
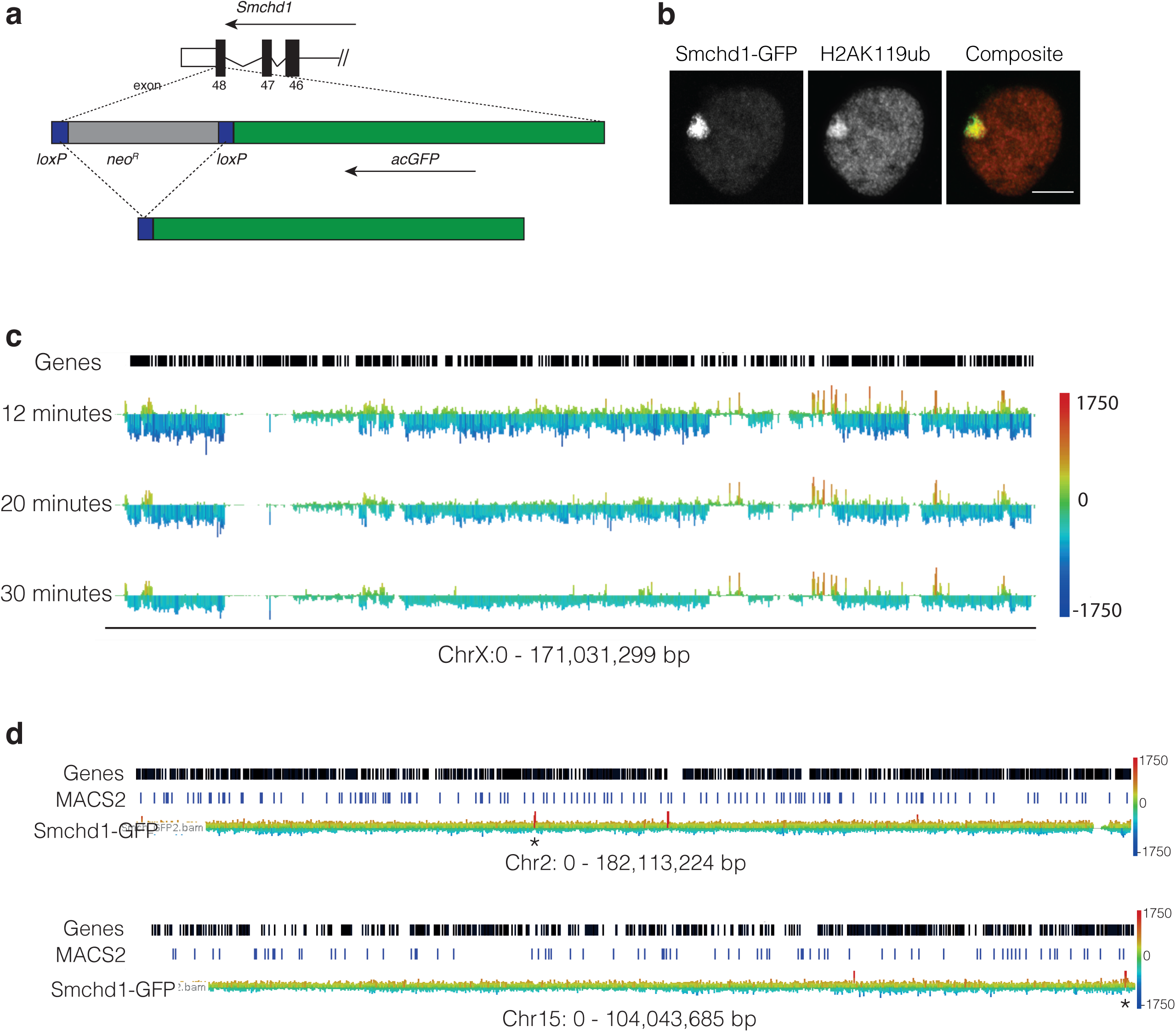
Generation of *Smchd1*^GFP^ knockin allele and its use for ChIP-seq. **a.** acGFP-loxP-neomycin resistance-loxP was knocked in immediately before the stop codon of *Smchd1* in 129T2/SvEms strain mESC, such that Smchd1 would be produced as a Smchd1-GFP fusion protein. Correctly targeted mESC clones were used to generate a line of mice, and the neomycin resistance cassette removed by Cre-mediated recombination. **b.** The Smchd1-GFP fusion protein was detected on the inactive X chromosome, as marked by H2aK119ub. GFP and H2AK119ub immunofluorescence performed in female *Smchd1*^GFP/GFP^ mouse embryonic fibroblasts. **c.** Optimisation of the ChIP-seq sonication conditions to release the Xi from the insoluble fraction. Sonication was performed in X_129_^XistΔA^X_Castaneus_ NSCs. Screenshot from genome browser showing the X chromosome. Positions of genes are shown as black boxes along the top. Tracks displayed are showing normalised counts for reads that mapped to the inactive X chromosome (castaneus) minus the reads that mapped to the active X chromosome (domesticus). Height and colour of bar represents relative coverage. Sonication time is labelled on the left. As sonication time increases, reads that map to the Xi increase. Note that even after 30 minutes sonication, coverage from the Xi is still not as great as that from the Xa (relative coverage across the X chromosome is negative). Scale bar on right is read count. Data from n=1. **d**. Enrichment of GFP over WCE on Chromosome 2 and Chromosome 15. Height and colour of peaks indicates strength of read counts normalised to WCE; red is most enriched, blue is most depleted. *HoxD* locus on Chromosome 2 and *HoxC* locus on Chromosome 15 are marked with an asterisk. Position of MACS2-like peaks called using Seqmonk are indicated in blue.

**Supplementary Figure 4.**
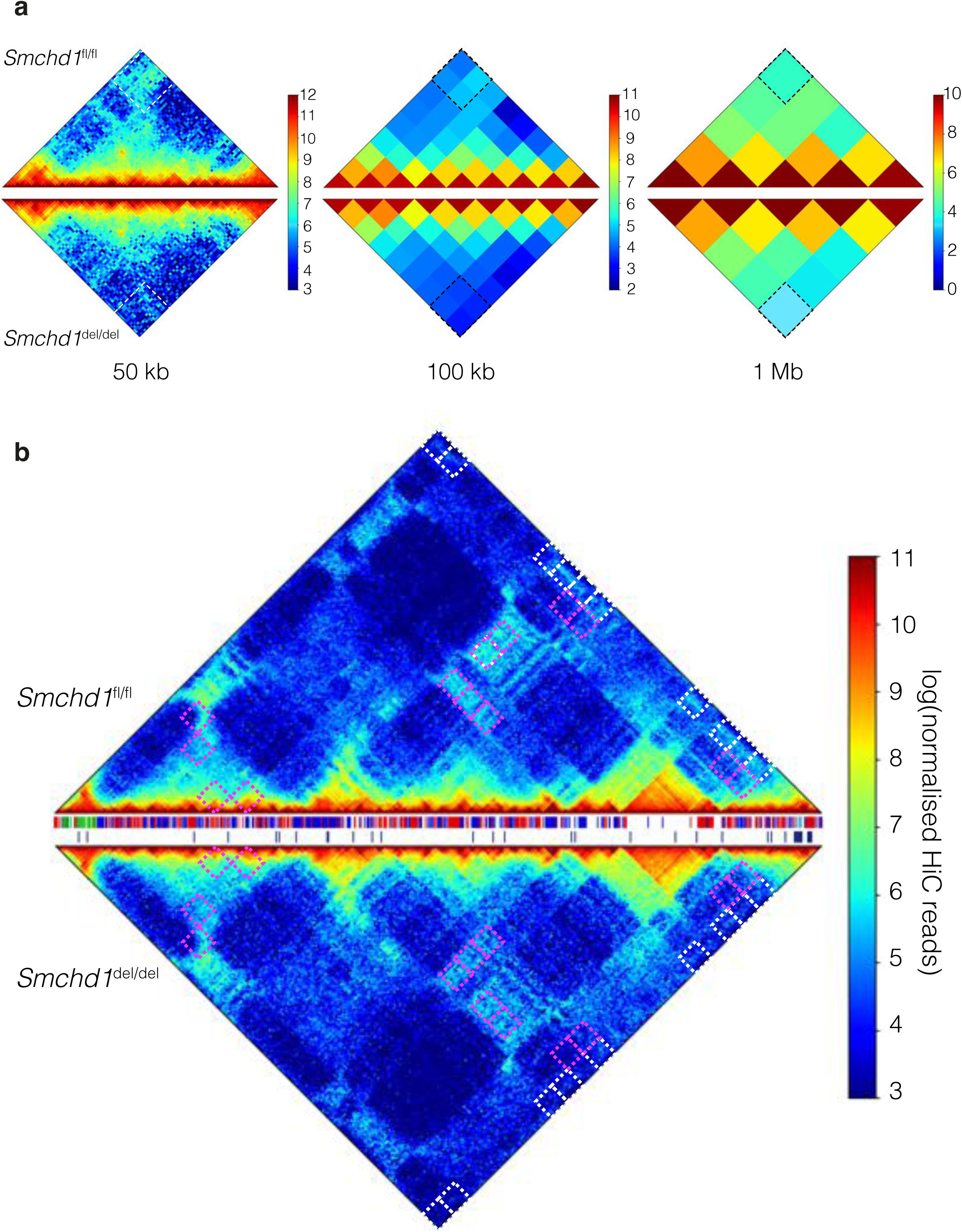
Normalised HiC interaction frequencies for interactions between the HoxB cluster and other clustered genes. **a.** Normalised Hi-C interaction frequencies at 50kb, 500 kb and 1Mb resolution displayed as a heat map and rotated 45^°^ for Chr11:96-100Mb containing the *HoxB* and *Keratin* gene clusters. Above the horizontal is the *Smchd1*^fl/fl^ heatmap, below the horizontal *Smchd1*^del/del^. The colour of the contact map, from blue to red, indicates the log_2_(contact frequency). Dotted white/black regions indicate the region of differential interaction, which is visible in the analyses at all 3 resolutions. **b.** Heatmaps displayed as in **a**, at 100kb resolution, for the region between the Olfactory receptor cluster genes (73 Mb) and *HoxB* cluster (97 Mb) on chromosome 11. Between the heatmaps, genes are shown as bars (green for Olfactory receptor genes), Smchd1 ChIP-seq peaks are shown as black bars. The white boxes bound the DIs with one anchor in the *HoxB* cluster. The magenta boxes are all other DIs in the region.

**Supplementary Figure 5.**
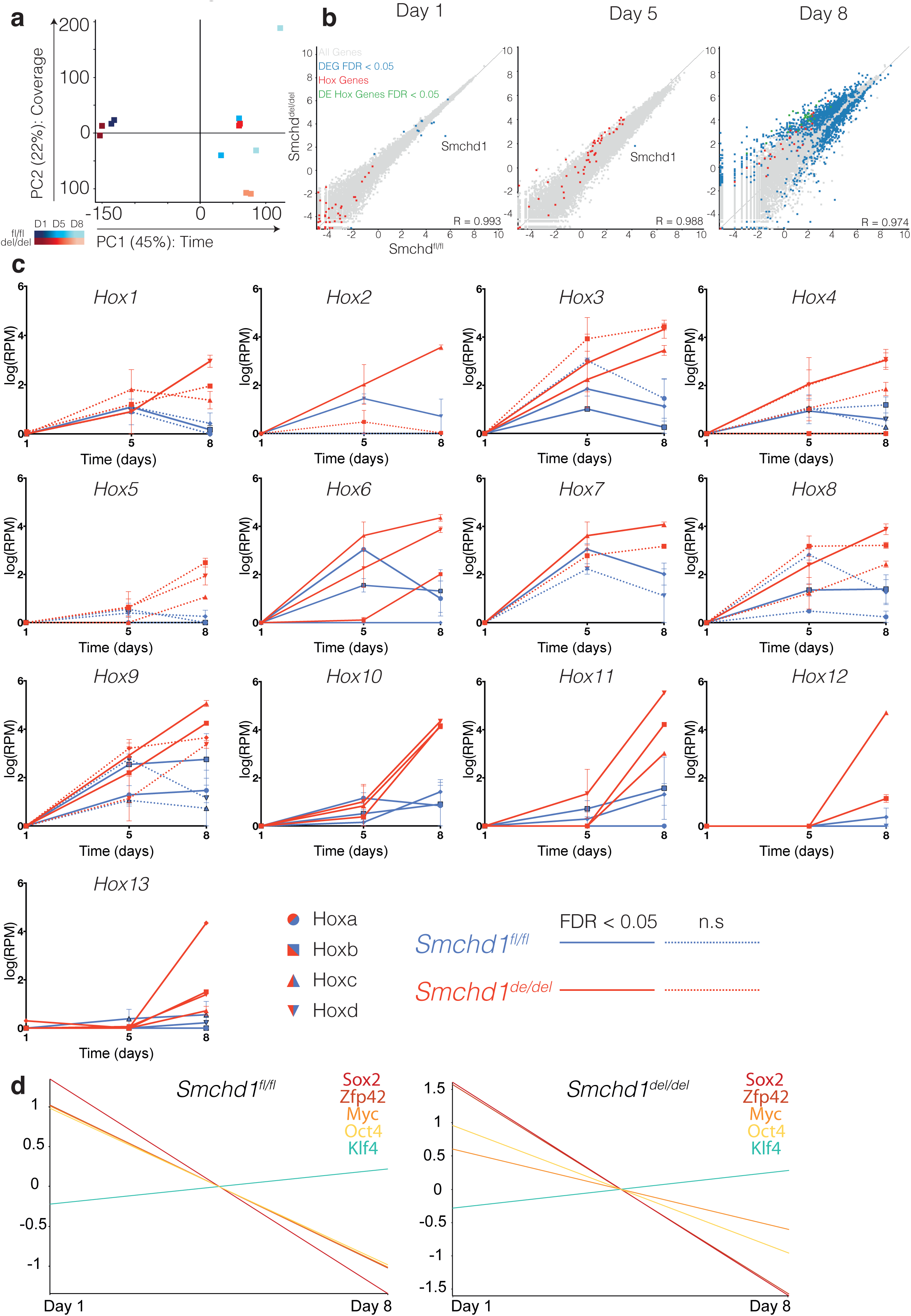
*Hox* genes are upregulated in *Smchd1* null NMPs differentiated from mESC. **a.** Principal component analysis (PCA) of RNAseq libraries from the differentiation of *Smchd1*^fl/fl^ and *Smchd1*^del/del^ mESCs to NMPs. Samples separate by timepoint on the first component (45%, x-axis) and by coverage on the second (22%, y-axis). *Smchd1*^fl/fl^ samples are depicted in shades of red, and *Smchd1*^del/del^ samples are depicted in shades of blue. Darker colours represent early time points, whereas lighter colours represent later time points. **b.** Dot plots showing expression of all genes in *Smchd1*^fl/fl^ (x-axis) compared with *Smchd1*^del/del^ (y-axis) in log(RPM) at D1, D5, and D8 post-induction of NMP differentiation. All genes are depicted in grey. Differentially expressed genes (FDR < 0.05) are depicted in blue. *Hox* genes are depicted in red. *Hox* genes which are differentially expressed are depicted in green. Pearson’s correlation value (R) is shown on the bottom, right of each scatter plot. **c.** Line graphs showing *Hox* gene expression throughout *in vitro* differentiation of male *Smchd1*^fl/fl^ and *Smchd1*^del/del^ mESCs into NMPs, as shown in Fig. 3a. Each graph shows expression for one paralogue group. i.e. the top left graph is showing expression of the *Hox1* gene in three paralogous clusters. Time in days (x axis) is plotted against log_2_ transformed reads per million (RPM) (y axis). *Smchd1*^fl/fl^ values are plotted in blue, and *Smchd1*^del/del^ in red. Circles represent genes from the *HoxA* cluster, squares represent *HoxB*, triangles represent *HoxC*, and upside-down triangle for those in *HoxD*. Solid lines depict genes that are differentially expressed (FDR < 0.05) at D8 between *Smchd1*^fl/fl^ and *Smchd1*^del/del^ NMP. Dotted lines indicate genes that were not called as differentially expressed using edgeR. **d.** Line graphs showing the downregulation of 4/5 pluripotency genes between day 1 and day 8 of the differentiation series in *Smchd1*^fl/fl^ and *Smchd1*^del/del^ samples.

**Supplementary Figure 6.**
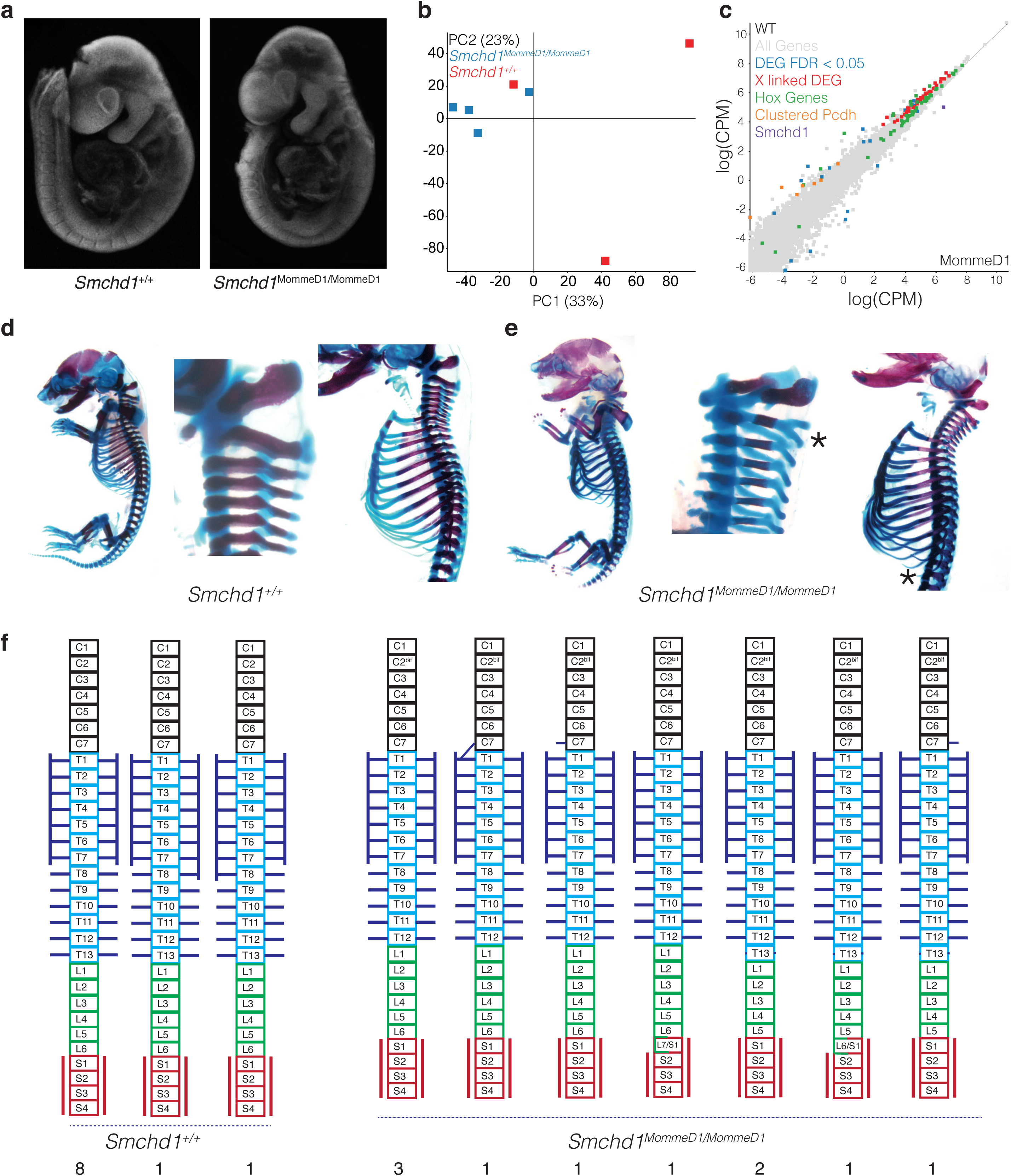
*Smchd1* null embryos have altered posterior Hox gene expression and skeletal abnormalities. **a.** Images of a representative somite-matched, sex-matched pair of E9.5 embryos, *Smchd1*^+/+^ and *Smchd1*^MommeD1/MommeD1^. **b.** Principal component analysis (PCA) of RNA-seq libraries from the somite-matched female paired *Smchd1*^+/+^ (n=3) and *Smchd1*^MommeD1/MommeD1^ (n=4) samples. *Smchd1*^+/+^ samples are depicted in green, and *Smchd1*^MommeD1/MommeD1^ samples are depicted in blue. **c.** Gene expression data from somite-matched female paired *Smchd1*^+/+^ (n=3) (y-axis) against *Smchd1*^MommeD1/MommeD1^ (n=4) samples (x-axis). Genes that are differentially expressed between *Smchd1*^+/+^ and *Smchd1*^MommeD1/MommeD1^ samples (FDR<0.05) are shown in blue. The *Hox* genes are shown in green, X-linked DEG in red, clustered protocadherins in orange, and Smchd1 in purple. All other genes are represented in grey. Axes are mean log transformed reads per million. **d-e.** Representative images from n = 10 whole mount skeletons of *Smchd1*^+/+^ (**d**) and *Smchd1*^MommeD1/MommeD1^ (**e**) E17.5 embryos, stained with Alizaren red and Alician blue. Whole body image, higher magnification view of the cervical vertebrae, and higher magnification sagittal view of embryo shown in d. *Smchd1*^MommeD1/MommeD1^ embryo shows abnormal bifurcating morphology of C2, and absence of T13 rib, indicated by asterisks. Note image for C2 is not the same *Smchd1*^MommeD1/MommeD1^ embryo as full body and sagittal view. **f.** Summarised data from scoring of whole mount skeletons from *Smchd1*^+/+^ (n=10) and *Smchd1*^MommeD1/MommeD1^ (n=10) male embryos. Cervical, thoracic, lumbar and sacral vertebrae are indicated, with ribs shown by horizonal blue lines, sternum shown by vertical blue lines and sacrum shown by vertical red lines. Dark blue dots indicate dramatically reduced rib process. C2^bif^ refers to a malformed C2 vertebra, where it bifurcates at the dorsal side.

**Supplementary Figure 7.**
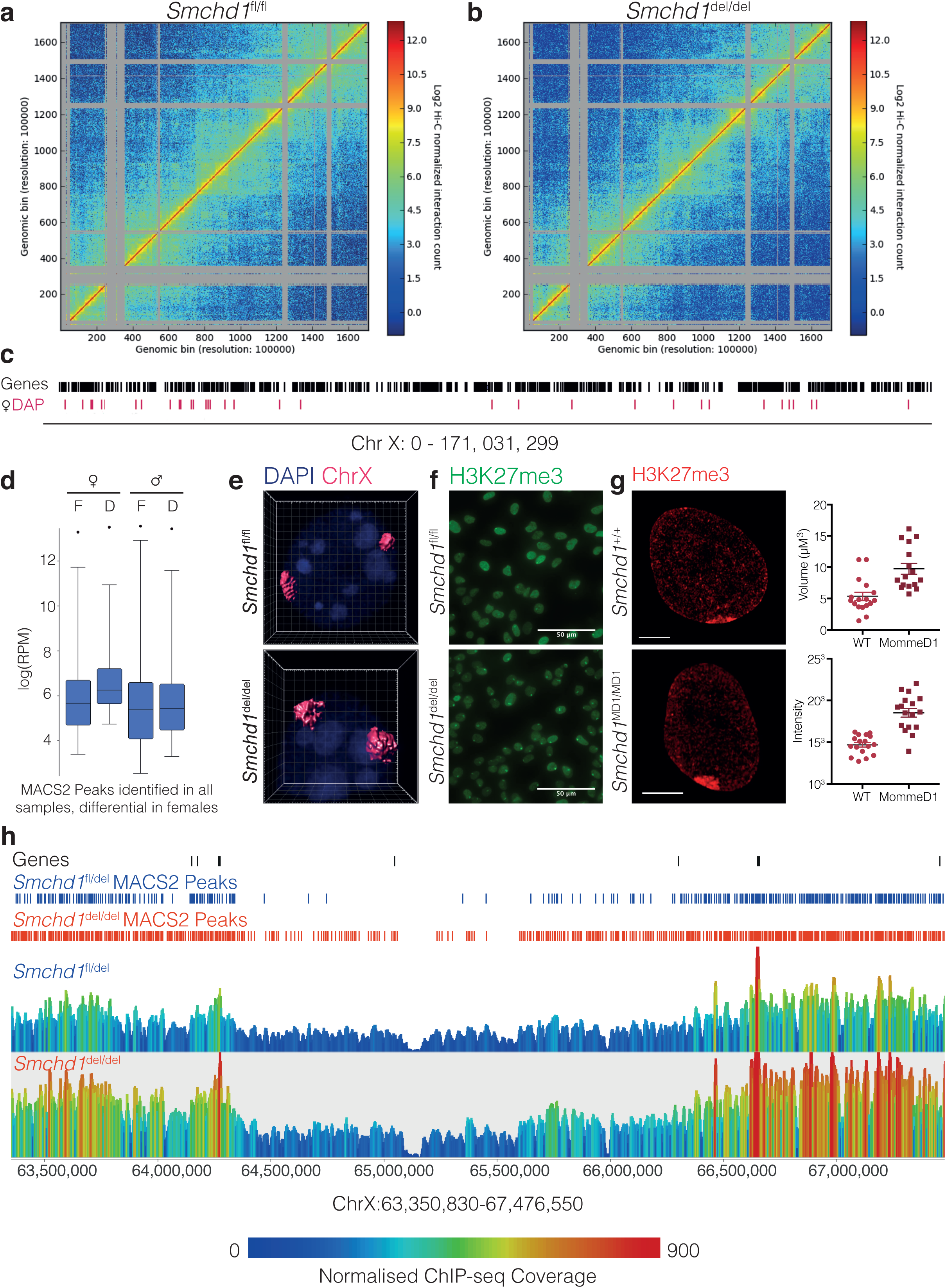
*Smchd1* deletion does not alter the compaction accessibility of the inactive X chromosome, but does result in enriched H3K27me3 on the Xi. **a.** Normalised Hi-C interaction frequencies at 100 kb resolution, generated from 200 million reads randomly subsampled from libraries at each condition, displayed as a heat map for the X chromosome in *Smchd1*^fl/fl^ (**a**) and *Smchd1*^del/del^ (**b**) NSCs. The colour of the contact map, from blue to red, indicates the log_2_(contact frequency). **c.** ATAC-seq was performed in *Smchd1*^fl/fl^ and *Smchd1*^del/del^ NSCs (n=3 female, n=2 male), one week post Cre mediated deletion of *Smchd1*. Position of female specific X-linked differentially accessible peaks (DAP) from the ATAC-seq analysis called using and edgeR analysis on MACS2 peaks in red. Positions of genes are indicated in black along the top. **d.** Sex-specific differential analysis of the MACS2 peaks identified in all samples, found 540 female-specific DAPS genome-wide. These were regions that became more accessible in the absence of Smchd1 in females, but not in males, as shown by box plots showing the coverage at 36 differential X-linked ATAC-seq peaks in log (RPM) between female *Smchd1*^fl/fl^ (F) and *Smchd1*^del/del^ (D) NSCs (left, n=3), and males (right, n=2). The female specific DAPs were not enriched on the X chromosome relative to autosomes, X-linked DAPs comprising 6% of the total female specific DAP (36/540). p values shown were determined by Student’s two-tailed t test. **e.** Confocal imaging of *Smchd1*^fl/fl^ (top) and *Smchd1*^del/del^ (bottom) NSC nuclei following DNA FISH with X chromosome paint, n=23 observations per genotype, n=2 cell lines. DAPI stained DNA is shown (blue) and X chromosome paint (magenta), shown with 3D rendering performed in Imaris to measure volume. **f.** Widefield images of H3K27me3 immunofluorescence performed in female *Smchd1*^fl/fl^ (top) and *Smchd1*^del/del^ (bottom) NSCs (green). **g**. Confocal images with Airyscan processing of H3K27me3 immunofluorescence (red) in female *Smchd1*^+/+^ (top) and *Smchd1*^MommeD1/MommeD1^ NSCs (bottom). Dot plots of volume and mean intensity of focal H3K27me3 enrichment in 3D using Imaris. Statistical comparisons were made using a two-tailed student’s t-test, line indicates mean, error bars are SEM. Measurements from 17 observations in n=2 cell lines, p <0.01. **h.** Genome browser tracks for the H3K27me3 ChIP-seq from ChrX:63,350,830-67,476,550 on the Castaneus-derived Xi chromosome in *Smchd1*^del/fl^ and *Smchd1*^del/del^ NSCs (n=2). Genes are shown in black, MACS2-called peaks in *Smchd1*^del/fl^ NSCs are shown in blue, and MACS2-called peaks in *Smchd1*^del/del^ NSCs are shown in red. Scale is normalised ChIP-seq read counts, normalised for read depth using the total autosomal read count.

## Supplementary Tables List

Supplementary Table 1. Differential interactions between *Smchd1*^fl/fl^ and *Smchd1*^del/del^ female NSCs, *Smchd1*^fl/fl^ and *Smchd1*^del/del^ male NSCs, based on diffHic analysis at 1 Mb, 500kb and 100 kb resolutions.

Supplementary Table 2. Strengthened and weakened TAD boundaries, TAD insulation scores, and compartment boundaries based on Tadbit analysis in *Smchd1*^fl/fl^ and *Smchd1*^del/del^ female NSCs.

Supplementary Table 3. Smchd1-GFP ChIP-seq peaks.

Supplementary Table 4. Loops defined by HiCCUPS in *Smchd1*^fl/fl^ and *Smchd1*^del/del^ female NSCs.

Supplementary Table 5. Differentially expressed genes between *Smchd1*^fl/fl^ and *Smchd1*^del/del^ female NSCs.

Supplementary Table 6. Differential ATAC-seq peaks between *Smchd1*^fl/fl^ and *Smchd1*^del/del^ NSCs.

Supplementary Table 7. H3K27ac and H3K27me3 ChIP-seq peaks from *Smchd1*^fl/fl^ and *Smchd1*^del/del^ female NSCs.

Supplementary Table 8. Differentially expressed genes between *Smchd1*^fl/fl^ and *Smchd1*^del/del^ male ES cells differentiating to neuromesodermal progenitors at days 1, 5 and 8 of differentiation.

Supplementary Table 9. Differentially expressed genes between presomitic mesoderm from *Smchd1*^+/+^ and *Smchd1*^MommeD1/MommeD1^ E9.5 female embryos.

Supplementary Table 10. Read counts for X-linked genes that mapped to the active X chromosome (genome 1) and the inactive X chromosome (genome 2) in *Smchd1*^del/fl^ and *Smchd1*^del/del^ female NSCs from RNAseq.

Supplementary Table 11. H3K27me3 ChIP-seq peaks on the Xi from *Smchd1*^del/fl^ and *Smchd1*^del/del^ female NSCs.

Supplementary Table 12. Read counts for RRBS data from the Xa and Xi of *Smchd1*^del/fl^ and *Smchd1*^del/del^ female NSCs

Supplementary Table 13. Oligonucleotides used for genotyping (within online methods file).

## Online methods

All genomics data can be found in the Gene Expression Omnibus, under GSE111726. To review this accession, go to https://www.ncbi.nlm.nih.gov/geo/query/acc.cgi?acc=GSE111726 Enter token crufsamaxrwbzml into the box.

### Mouse strains

All mice were bred and maintained using standard animal husbandry procedures, approved by the WEHI Animal Ethics Committee under animal ethics numbers AEC 2014.026 and AEC 2018.004.

*Smchd1*^fl/fl^ mice were generated on a C57BL/6 background, as described previously^1^. Mice carrying the *Smchd1*^MommeD1^ mutation as previously described^2^ were maintained on the FVB/N inbred background. Both strains were genotyped as previously described (primers in Supplementary Table 13). *Xist*^*Δ*A^ mice were generated on a 129 background as previously described^3^. Castaneus strain (Cast/EiJ) animals were obtain from Jackson laboratories.

The *Smchd1*^GFP^ targeted allele was constructed by standard gene targeting techniques using the mouse C1368 ES cell line (129T2/SvEms strain). The 5’ (chr17:71,345,341-71,347,443) and 3’ (chr17:71,343,347-71,345,340) homology regions for the targeting construct were generated by Pfu polymerase high fidelity PCR. These homology regions were cloned either side of an acGFP ORF (Clontech) and a loxP flanked neo selection cassette so that the acGFP-loxP-neo-loxP cassette was introduced in-frame immediately before the Smchd1 stop codon. The engineered *Smchd1*^GFP^ allele was designed to produce a fusion protein consisting of wild type Smchd1 with a carboxy terminal acGFP moiety. We specifically chose acGFP rather than EGFP because acGFP is known not to dimerise. The targeting construct was introduced into the ES cell line by electroporation. G418 surviving cell clones were screened by Southern blotting of XbaI cut genomic DNA to identify correctly targeted clones. The probe used for Southern blotting was generated by PCR amplification of an upstream genomic region (chr17:71,348,361-71,349,085) that was external to the targeting homology arms but within an XbaI fragment (generated by the XbaI sites at chr17:71,344,994-71,344,999 and chr17:71,349,766-71,349,771) that could be used as diagnostic for correctly targeted clones. Since the targeting construct introduces an XbaI site into the genome, correctly targeted clones were identified by a shift of the usual 4772bp band to a 6691bp band. Correctly targeted ES cell clones were then used for mouse chimaera generation by injection of the ES cells into C57Bl6/J strain blastocysts. Once chimaeras were identified and germline transmission established, the loxP flanked neo selection cassette was removed by crossing the targeted mice with Cre-deleter mice (E2A-Cre). Crossing of the resulting *Smchd1*^GFP^ allele carrying mice to homozygosity indicated that the Smchd1-acGFP fusion protein was likely to be fully functional since female *Smchd1*^GFP/GFP^ mice were obtained at similar frequencies to *Smchd1*^+/+^ (wild-type) female animals – unlike the loss of function *Smchd1*^MommeD1^ allele which is female embryonic lethal when homozygous. Smchd1-GFP localised to the inactive X as has been shown for wild-type Smchd1 protein. The *Smchd1*^GFP^ allele was backcrossed onto the C57BL/6J strain for 10 generations then maintained as a homozygous congenic line of mice. This strain was genotyped by PCR using oligos specific to the integration site (Supplementary Table 13).

### Embryo genotyping

Embryo tails or yolk sacs were used to prepare DNA using standard methods. Genotypes were determined by PCR using GoTaq Green (Promega) or allelic discrimination for MommeD1. Primers for genotyping were as for standard genotyping of the strain, except that sex was additionally determined by PCR for *Otc* (X chromosome) and *Zfy* (Y chromosome) (Supplementary Table 13). Sex was often also determined at E14.5 by dissection of embryonic gonads.

### Derivation of Mouse Embryonic Stem Cells (mESCs)

Female mice were super-ovulated by injecting 5 IU folligon (MSD Animal Health Australia) two days prior to, and 5 IU chorulon (MSD Animal Health Australia) on the day of mating. At E3.5, dams were sacrificed, uteri removed and blastocysts flushed from the uterine horns with M2 medium (Sigma-Aldrich). Blastocysts were washed in M2 medium twice, and 2i+LIF medium [KnockOut DMEM (Life Technologies), 1X Glutamax (Life Technologies), 1 X MEM Non-Essential Amino Acids (Life Technologies), 1 X N2 Supplement (Life Technologies), 1 X B27 Supplement (Life Technologies), 1 × 2-mercaptoethanol (Life Technologies), 100 U/mL Penicillin/100 *μ*g/mL Streptomycin (Life Technologies), 10 *μ*g/mL Piperacillin (Sigma-Aldrich), 10 *μ*g/mL Ciprofloxacin (Sigma-Aldrich), 25 *μ*g/mL Fluconazol (Selleckchem), 1000 U/mL ESGRO Leukemia Inhibitory Factor (Merck), 1 *μ*M StemMACS PD0325901 (Miltenyi Biotech), 3 *μ*M StemMACS CHIR99021 (Mitenyi Biotech)] twice. Blastocysts were plated in non-tissue culture treated 24-well plates in 2i+LIF medium. Following 7 days in cultured at 37°C in a humidified atmosphere with 5 % (v/v) carbon dioxide and 5 % (v/v) oxygen, outgrowths were moved by mouth-pipetting through trypsin-EDTA for 2 minutes, mESC wash media [KnockOut DMEM (Life Technologies), 10 % KnockOut Serum Replacement (Life Technologies), 100 IU/ml penicillin/100 *μ*g/ml streptomycin (Life Technologies)], and finally 2i+LIF. Outgrowths were disrupted by pipetting, and transferred into a 24-well plate to be cultured as mESC lines.

mESCs were maintained in suspension culture in serum-free 2i+LIF medium on non-tissue culture treated plates at 37°C in a humidified atmosphere with 5 % (v/v) carbon dioxide and 5 % (v/v) oxygen. mESC were passaged daily. mESC colonies were collected and allowed to settle in a tube for < 5 minutes. Supernatant containing cellular debris was removed, and mESC colonies were resuspended in Accutase (Sigma-Aldrich) and incubated at 37°C for 5 minutes to achieve a single-cell suspension. At least 2X volume of mESC wash media was added to the suspension, and cells were pelleted by centrifugation at 600 × g for 5 minutes.

### Differentiation of mESC into Neuromesodermal Progenitors (NMPs)

This protocol was adapted from three previous reports^4-7^. Four days prior to differentiation, mESCs were put onto tissue-culture ware that had been coated with 0.1% porcine gelatin (Sigma-Aldrich). mESC were passaged into 75 % 2i+LIF medium 25 % ES cell DMEM+LIF [High glucose DMEM, 0.085 mM MEM Non-Essential Amino Acids (Life Technologies), 34 mM NaHCO3, 0.085 mM 2-mercaptoethanol (Life Technologies), 100 *μ*g/ml streptomycin, 100 IU/ml penicillin, 15% foetal bovine serum (FBS) (Life Technologies), 1000 U/mL ESGRO Leukemia Inhibitory Factor (Merck), 10 *μ*g/mL Piperacillin (Sigma-Aldrich), 10 *μ*g/mL Ciprofloxacin (Sigma-Aldrich), 25 *μ*g/mL Fluconazol (Selleckchem)]. After 24 h, the medium was changed to 50 % 2i+LIF medium 50 % ES cell DMEM+LIF. After 48 h, the medium was changed to 25 % 2i+LIF medium 75 % ES cell DMEM+LIF. After 72 h, the cells were split and seeded at 2.5 × 10^5^ cells per cm^2^ in 100 % ES cell DMEM+LIF. The next morning (D0 differentiation), ES cell DMEM+LIF was removed, cells were washed with PBS to remove any traces of FBS, and medium was replaced with N2B27 medium [1:1 mix of Advanced DMEM/F12 (Life Technologies) and Neurobasal media (Life Technologies), 0.5 X N2-supplement (Life Technologies), 0.5 X B27-supplement, 1 X Glutamax (Life Technologies), 40 *μ*g/mL BSA Fraction V (Life Technologies), 1X Beta-mercaptoethanol (Life Technologies), 100 U/mL Penicillin/100*μ*g/mL Streptomycin (Life Technologies), 10 Piperacillin *μ*g/mL (Sigma-Aldrich), 10 *μ*g/mL Ciprofloxacin (Sigma-Aldrich), 25 *μ*g/mL Fluconazol (Selleckchem)] supplemented with 10 ng/mL recombinant human basic FGF (Peprotech). After 48 hours (D2), medium was replaced with N2B27 medium supplemented with 10 ng/mL recombinant human basic FGF (Peprotech) and 5 *μ*M StemMACS CHIR99021 (Mitenyi Biotech). After a further 48 hours (D4), medium was replaced with N2B27 plus bFGF and StemMACS CHIR99021. After 48 hours (D6), medium was replaced with N2B27 medium supplemented bFGF, StemMACS CHIR99021, and 50 ng/mL recombinant human GDF11 (Mitenyi Biotech). NMPs were maintained in N2B27 medium supplemented bFGF, StemMACS CHIR99021 and GDF11.

### Generation of Neural Stem Cells (NSCs)

NSCs were derived as exactly previously described^8^ and cultured with minimal changes. For culture, cells were seeded onto either plates coated with 15 ng/mL polyornithine (Sigma-Aldrich) and 10 ng/mL laminin (Sigma-Aldrich), or directly into tissue culture treated plates with 10 ng/mL laminin (Sigma-Aldrich) added to the medium, at a density of 40,000 cells per cm^2^. Cells were cultured at 37°C in 5% (vol/vol) CO_2_ incubator and passaged every 2 days using Accutase (Sigma–Aldrich) to detach the cells from the plates. Primary cells were maintained for a maximum of 20 passages.

### Generation of *Smchd1*^MommeD1/MommeD1^ MEFs

MEFs were generated as described from E10.5 *Smchd1*^MommeD1/MommeD1^ embryos^9^, and cultured at 37°C in 5% CO_2_ and 5% O_2_ for one week. Cells were passaged using 0.5% trypsin-EDTA (Life Technologies) and seeded at approximately 40,000 cells/cm^2^ on gelatin coated cover slips for immunofluorescence.

### Generation of *Smchd1*^del/del^ NSCs

*Smchd1*^fl/fl^ NSCs were transduced as described below with concentrated retroviral supernatant containing the MSCV-Cre-Puromycin construct. The next day, cells were selected with 1 *μ*g/mL Puromycin (Sigma-Aldrich) for at least 7 days. *Smchd1* deletion was confirmed by PCR, RTqPCR, and Western Blot.

### Generation of *Smchd1*^del/del^ mESCs

*Smchd1*^fl/fl^ NSCs were transduced as described below with concentrated retroviral supernatant containing either the MSCV-Cre-Puromycin or MSCV-Puromycin construct. The next day, cells were selected with 1 *μ*g/mL puromycin (Sigma-Aldrich) for at least 7 days. Smchd1 deletion was confirmed by PCR and RNA-seq. These NSCs do not contain polymorphic SNPs between each X chromosome and the genome as the *Smchd1*^fl/fl^ mouse line is inbred C57BL/6

X_129_^XistΔA^X_Castaneus_ *Smchd1*^del/del^ NSCs were generated by Cre-mediated deletion as above from X_129_^XistΔA^X_Castaneus_ *Smchd1*^del/fl^ NSCs. X_129_^XistΔA^X_Castaneus_ *Smchd1*^del/+^ controls were generated by Cre-mediated deletion from X_129_^XistΔA^X_Castaneus_ *Smchd1*^fl/+^ NSCs.

### Retrovirus Production and Transduction

VSVg pseudotyped retroviral supernatants were produced using calcium phosphate mediated transient transfection of 293T cells, as previously described^10^. The media was collected at 48 and 72 hours post-transfection, centrifuged to remove residual 293T cells, and either snap frozen or concentrated using Poly(ethylene glycol).

For transduction of NSCs, NSCs were seeded at a density of 40,000 cells per cm^2^ 8-16 hours before transduction. PEG concentrated viral supernatant was added to the culture together with polybrene at a final concentration of 4 *μ*g/mL. On the next day, cells were selected with 1 *μ*g/mL puromycin.

For transduction of mESCs, ESCs were seeded at 10^5^ cells per cm^2^ on plates that had been coated with 0.1% gelatin or in suspension at 10^5^ cells per mL in 2i+LIF medium containing PEG concentrated viral supernatant and 8 *μ*g/mL polybrene. The next day medium was changed and cells were selected with 1 *μ*g/mL puromycin.

### Dissection of the presomitic mesoderm from E9.5 embryos

The PSM was dissected from *Smchd1*^MommeD1/MommeD1^ *Smchd1*^MommeD1/+^ and *Smchd1*^+/+^ E9.5 embryos as previously described^11^. In brief, under a light microscope the tail bud was removed approximately 1.5 somite lengths after final formed somite. The PSM was snap frozen for later RNA harvest using a Qiagen RNeasy micro kit. The yolk sac was taken for genotyping. The embryo body was fixed in 4% paraformaldehyde for 2 hours, then washed through a methanol series as previously described^11^. To image, the embryo was dipped in ethidium bromide solution then photographed on a fluorescence dissection microscope to count somites. RNA-seq comparisons were made between somite-matched, sex-matched *Smchd1*^MommeD1/MommeD1^ and *Smchd1*^+/+^ PSM samples.

### Immunofluorescence

Immunofluorescence was performed on *Smchd1*^GFP/GFP^, *Smchd1*^fl/fl^ and Smchd1^del/del^ NSCs and *Smchd1*^+/+^, *Smchd1*^MommeD1/MommeD1^ MEFs as described in Chaumeil *et al*.^12^, with modifications. In brief, cells were seeded at 70% confluence the night before the experiment on polyornithine and laminin coated chamber slides. The next day, cells were washed in PBS, and fixed in 3% (w/v) paraformaldehyde made in PBS for 10 minutes at room temperature. Cells were washed three times in PBS for 5 minutes each, and then permeabilised in cold 0.5 % (v/v) Triton X-100 in PBS on ice for 5 minutes. Cells were washed three times in PBS for 5 minutes each, and then blocked in 1% (w/v) Bovine Serum Albumin (BSA) (Life Technologies) for at least 15 minutes. Cells were then incubated with a primary antibody diluted in 1% (w/v) BSA for 45 minutes at room temperature in a humid chamber. Cells were washed three times in PBS for 5 minutes each, and then incubated with a secondary antibody conjugated to a fluorophore diluted in 1% (w/v) BSA for 40 minutes at room temperature in a dark and humid chamber. Cells were washed three times in PBS for 5 minutes each, and were mounted in Vectashield HardSet mounting medium with DAPI (Vector Laboratories). When *Smchd1*^GFP/GFP^ cells were used, detection of Smchd1 protein was enabled via detection of native Smchd1-GFP fusion protein. Cells were visualised on an Elite Widefield (DeltaVision), LSM 880 (Zeiss), or Live-Cell AxioObserver (Zeiss) microscope. Images were analysed using the open source ImageJ distribution package, FIJI^13^. For each experiment 100–250 nuclei were analysed.

Imaris Software (Bitplane) was used to measure the volume occupied by markers of the Xi (H3K27me3 or X chromosome paint) within the nucleus. A threshold was manually set to measure the signal only above nucleoplasmic staining or background. A region of interest was then defined based on DNA paint or focal H3K27me3 enrichment. Volume was then calculated for the region of interest above the threshold.

### DNA FISH

DNA FISH was performed on X_129_^XistΔA^X_Castaneus_ *Smchd1*^del/del^ and *Smchd1*^+/fl^ NSCs. 1 *μ*g RP23-196F5 (*HoxB* locus) or RP2390M1 (*Keratin* locus) BAC DNA (CHORI) were used in a nick translation reaction (Vysis) to generate DNA probes labelled with SpectrumRed dUTP or SpectrumGreen dUTP (Vysis), respectively. ∼100 ng probe per sample was precipitated in Ethanol with 10% NaOAc, and 1 *μ*g Salmon Sperm (Life Technologies), and 1*μ*g Mouse Cot-1 DNA before being resuspended in formamide (Sigma-Aldrich), denatured at 75°C for 10 minutes, and allowed to compete with Cot-1 DNA for 1 hour at 37°C. Cells were prepared for DNA FISH exactly as described in Chaumeil *et al.*^12^. Cells were mounted in Antifade Mounting Medium (Vectashield) and visualised on an LSM 880 (Zeiss) microscope, with Airyscan processing. Images were analysed using the open source ImageJ distribution package, FIJI. Images were processed with the Median 3D filter to reduce background signal, and each image was manually thresholded with the Yen thresholding method. Distances between FISH signals were measured using the ImageJ plugin, DiAna^14^. Overlapping or touching FISH signals for *HoxB* and *Keratin* probes were scored as interacting, whereas those where paired probes fell within 4*μ*m but were not touching were scored as not interacting. 166 alleles were scored for the *Smchd1*^fl/+^ NSCs, 152 alleles were scored for the *Smchd1*^del/del^ NSCs.

### X Chromosome Paint

Female *Smchd1*^fl/fl^ and Smchd1^del/del^ NSCs were harvested with Accutase (Sigma-Aldrich), fixed for 5 minutes in Carnoys’ fixative (3:1 methanol:acetic acid), followed by centrifugation for 10 minutes at 4°C. Fixation and centrifugation were repeated, and nuclei were spotted onto a Superfrost Plus slide (Thermo Scientific) and allowed to dry. FITC-conjugated X-chromosome paint (AMP 0XG) was used as per the manufacturer’s instructions (Cytocell Ltd.). Cells were visualised on an LSM 880 (Zeiss) microscope, with Airyscan processing. Images were analysed using Imaris Software (Bitplane) as described above. For each experiment 10-20 nuclei were analysed.

### In *situ* Hi-C

*In situ* HiC was performed as described with modifications^15^. Briefly, 10^7^ *Smchd1*^fl/fl^ and Smchd1^del/del^ NSCs were cross-linked in 1% formaldehyde in NSC basal medium for 10 minutes at room temperature with rotation. Formaldehyde quenched with 200 mM Glycine, cells were washed with PBS, and then pellet was snap-frozen, and stored at −80°C. Nuclei were lysed in cold lysis buffer [10mM Tris-HCl pH8.0, 10mM NaCl, 0.2% Igepal CA630, 1 × Protease Inhibitor (Sigma)] and centrifuged at 600 × g for 5 minutes. Nuclei were resuspended in 0.5% (w/v) SDS and incubated at 62°C for 5 minutes. SDS was quenched with 10% Triton X-100 for 15 minutes at 37°C. Chromatin was digested by adding 200 U MboI with 200 mg/mL BSA for 1 hour, and a further 100 U every hour for 4 hours, and then incubated overnight at 37°C with rotation. MboI was inactivated by incubation at 62°C for 20 minutes. Restriction fragment overhangs were filled and marked with biotin by the addition of 300 *μ*M of each biotin-14-dATP (Life Technologies), dCTP, dGTP, and dTTP with 40 U DNA Polymerase I, Large (Klenow) Fragment (New England Biolabs), and incubation at 37°C for 45 minutes with rotation. Ligation buffer [1 × T4 DNA ligase buffer (New England Biolabs), 0.8% (v/v) Triton X-100, 1 mg/mL BSA (New England Biolabs)] and 2000 U T4 DNA Ligase (New England Biolabs) were added, and nuclei were incubated at room temperature with rotation for 4 hours. Nuclei were pelleted at 1000 × g for 5 minutes, resuspended in ligation buffer containing 1 mg/mL proteinase K (Roche) and 1% (w/v) SDS, and incubated at 55°C for 30 minutes. Crosslinks were reversed by the addition of 500 mM NaCl overnight incubation at 68°C. DNA was isolated by ethanol precipitation, and then dissolved in 10 mM Tris-HCl, pH 8. DNA was fragmented by sonication using the Covaris S220 sonicator (peak power: 175, duty factor: 10, cycle/burst: 200, duration: 58 s). Fragmented DNA was size selected by incubation with 0.55 × volume AMPure XP magnetic beads, and resuspended in 10 mM Tris-HCl, pH 8. Biotinylated DNA was enriched by incubation with Dynabeads MyOne Streptavidin T1 beads (Life Technologies) as per manufacturer’s instructions. Following two washes in tween wash buffer (TWB) [5mM Tris-HCl (pH 7.5), 0.5 mM EDTA, 1M NaCl, 0.05% Tween 20] at 55°C with mixing for 2 minutes each, DNA bound to the beads was resuspended in 1 × T4 DNA Ligase buffer with 10 mM ATP (New England Biolabs), 2 mM dNTP (New England Biolabs), 50 U T4 PNK (New England Biolabs), 12 U T4 DNA polymerase I (New England Biolabs) and 5 U DNA polymerase I, Large (Klenow) Fragment (New England Biolabs) and incubated at room temperature for 30 minutes to add a phosphate group for subsequent ligation steps, and to remove biotin from unligated ends. The beads were washed twice in TWB at 55°C with mixing for 2 minutes each, and A-tailing was performed by an on bead incubation of DNA in 1 × NEBuffer 2 (New England Biolabs), 500 *μ*M dATP (New England Biolabs), and 25 U Klenow exo minus (New England Biolabs) at 37°C for 30 minutes. Following two washes in TWB at 55°C with mixing for 2 minutes each, beads were washed in Quick ligation reaction buffer (New England Biolabs). 2 *μ*L DNA Quick ligase (New England Biolabs) was added to beads resuspended in 50 *μ*L Quick ligation reaction buffer (New England Biolabs), along with an Illumina indexed adapter, and incubated at room temperature for 15 minutes. Beads were washed twice in TWB at 55°C with mixing for 2 minutes each, and DNA was eluted by boiling beads in 10 mM Tris-HCl, pH 8 for 10 minutes. Libraries were amplified in 50 *μ*L Phusion PCR Master Mix [1 × High Fidelity Phusion Buffer (New England Biolabs), 200 *μ*M dNTPs (New England Biolabs), 5 *μ*L PCR Primer Cocktail (Illumina), 0.5 *μ*L Phusion Taq Polymerase (New England Biolabs)] with the following cycle conditions: 98°C for 30 seconds; 9 cycles of 98°C for 10 seconds, 60°C for 30 seconds, 72°C for 30 seconds; and 72°C for 5 minutes. DNA was isolated using AMPure XP magnetic beads, and libraries were subject to 75 bp paired-end sequencing on the Illumina NextSeq platform. Libraries were sequenced to a depth of at least 50 million reads (*Smchd1*^del/del^: 124860050; 100099110; 57467204. *Smchd1*^fl/fl^: 94218954; 200738168; 119847878).

Primary data processing was performed using HiCUP ^16^ v0.5.8. Differential Interactions were identified using diffHiC v1.12.0, which utilises edgeR statistics^17^. In brief, reads mapped and filtered using HiCUP were counted into 100 kb and 1 Mb bin pairs. Noise was removed by filtering out low abundance reads based on a negative binomial distribution and using inter-chromosomal counts to determine non-specific ligation events. Libraries were then normalised using TMM normalisation, and trended biases removed by fitting libraries to a generalised linear model. EdgeR was then used to test for differential interactions between genotypes, at either 100 kb or 1 Mb resolution, using a quasi-likelihood F-test, then we adjusted for multiple testing using FDR. Overlap with Smchd1 ChIP-seq peaks was performed using genomic coordinates for autosomal and X-linked MACS2 called ChIP peaks from GFP ChIP performed using MNase and sonication in *Smchd1*^GFP/GFP^ NSCs, respectively^18^.

### Generation of Hi-C contact matrices

To construct the Hi-C interaction matrices of the Smchd1 replete and deleted genomes, all reads were mapped to the *mm10* reference genome using TADbit^19^ v0.2.0.58 with the original iterative mapping strategy ICE^20^. TADbit uses the GEM mapper^21^ to map Fastq files. The minimal size for mapping was set to 25bp and the iterative mapping procedure increased in steps of 5bp until a maximal read length of 75bp was reached. A filtering strategy was then applied to correct for experimental biases as described in Imakaev *et al.*^19^. Once filtered, the read-pairs were binned at a 100 kb, 500 kb or 1 Mb resolution and columns containing few interactions counts were removed following the two-step strategy described in Serra *et al.*^19^. The remaining bins were further normalised using ICE^19^ as implemented in TADbit. As the resulting interaction matrices of the three separate Hi-C experiments for both conditions were highly correlated (Spearman correlation > 0.96), we merged the input reads into a single dataset, one for each condition. The new datasets were also filtered and normalised by TADbit and the resulting merged interaction maps were used to describe the 3D organisation of the *Smchd1* deleted and Smchd1 wild-type genome.

The subsampled contact matrices were constructed from a random selection of 200 million readpairs for both conditions respectively, then binned at a 100 kb resolution and then filtered and normalised following the same steps described previously.

### TAD detection on Hi-C contact maps

The merged interaction maps were used for domain detection at a 100kb resolution. The TADbit program^19,22^ employs a breakpoint detection algorithm that returns the optimal segmentation of the chromosome using a BIC-penalized likelihood^23^.

### Alignment of TAD boundaries

To assess whether TAD borders are conserved throughout different experiments, TADbit performed a multiple-experiment border alignment algorithm, starting from different border definitions of the same genomic region and aligns each TAD to a consensus TAD list, using the classic Needleman-Wunsch algorithm^24^.

### Compartment Detection on Hi-C Contact Maps

The merged interaction maps were used for compartments detection at a 1 Mb resolution. The TADbit program^19,22^ was used to search for compartments in each chromosome, by computing a correlation matrix on the normalised matrix and by using the first eigenvector to identify compartments.

### Annotation of chromatin looping with HiCCUPS

Annotation of chromatin looping was performed using the HiCCUPS peak calling algorithm^25^. Replicate HiC fastq files were merged for each condition and processed using the Juicer pipeline v1.8.9 to map HiC reads to the mm10 reference genome, remove duplicate and near duplicate reads, and reads that map to the same restriction fragment. HiC contact maps were produced from filtered reads with a MAPQ >30. HiCCUPS was run using default parameters, and identified 3192 loops in *Smchd1*^del/del^ HiC heatmaps, and 2410 loops in *Smchd1*^fl/fl^ HiC heatmaps.

### Transformed contact matrices

The merged interaction maps were used to compute the absolute distance and the mean distance between interacting bins and to construct the transformed interaction matrices. For each interaction m*_i,j_* defined as an interaction between the bin *i* and bin *j*, the absolute distance *AD* was computed as the absolute difference between the genomic location of the bin *i* and the bin *j*. The mean distance *MD* was computed as the mean between the genomic location of the bin *i* and bin *j*.

For an interaction m*_17,26_* of the merged interaction map at a 100 kb resolution for the X chromosome in *Smchd1*^fl/fl^

*AD* = |17 ∗ 100000 − 26 ∗ 100000| = 900000

*MD*= (17 ∗ 100000 + 26 ∗ 100000/2) = 2150000.

The differential plot was computed as the difference between the two transformed interaction matrices *Smchd1*^del/del^ versus in *Smchd1*^fl/fl^.

### RNA-seq

RNA was extracted using and RNeasy Mini or Micro kit (Qiagen) or a Quick RNA kit (Zymo). RNA was quantified using Qubit RNA Assay Kit (Life Technologies). Libraries were prepared using TruSeq RNA sample preparation kit from 1 *μ*g total RNA as per manufacturers’ instructions. 200-400bp products were size selected and cleaned up using AMPure XP magnetic beads. Final cDNA libraries were quantified using a Qubit dsDNA Assay Kit (Life Technologies) for sequencing on the Illumina NextSeq platform. C57BL/6 inbred NSC RNA-seq, ES cell RNA-seq and PSM RNA-seq libraries were subject to 75 bp single-end reads. 129/Cast NSC samples were subject to 75 bp paired-end reads. Reads were mapped with Tophat v2.0.12 using default options^26^. RNAseq data were analysed using Seqmonk v1.36.1 or v1.38.2. Reads under mRNA were quantified, and EdgeR statistics were used to test for differentially expressed genes in Seqmonk (FDR < 0.05)^27^. Geneset enrichment test for differentially expressed genes in *Smchd1*^MommeD1/MommeD1^ was performed using rotation gene set test (ROAST)^28^.

### ChIP-seq

ChIP for Smchd1 on the Xi was performed as described with modifications^29^. In brief, 4 × 10^7^ female *Smchd1*^GFP/GFP^ and *Smchd1*^+/+^ NSCs were harvested using Accutase (Sigma-Aldrich), washed in PBS, and were cross-linked with 1% formaldehyde (vol/vol) for 10 min at room temperature with rotation, then quenched with 125 mM Glycine. Cells were washed twice in ice-cold PBS, and pelleted by centrifugation at 600 × g at 4°C for 5 min. Approximately 2x 10^6^ were lysed with 1 mL of immunoprecipitation (IP) buffer [150 mM NaCl, 50 mM Tris·HCl (pH 7.5), 5 mM EDTA (pH 7.5), 0.5% (vol/vol) Nonidet P-40, 1.0% (vol/vol) Triton X-100] supplemented with 1 X c0mplete protease inhibitor (Roche). Nuclei were collected by centrifugation at 12,000 × × g at 4 °C for 1 min. The nuclei pellet was washed twice with IP buffer and then subjected to sonication in 130 *μ*L of IP buffer using the Covaris S220 sonicator (peak power: 125, duty factor: 10, cycle/burst: 200, duration: 1800 s). The chromatin was cleared, and the soluble chromatin was incubated with 10 *μ*g anti-GFP antibody (Life Technologies, A-11122) at 4°C overnight. Antibody-enriched fractions were captured by incubation with 20 *μ*L of Dynabeads Protein G (Life Technologies) at 4°C for 1, followed by 6X washes in IP buffer. Immunoprecipitated fractions were eluted in 1% (wt/vol) SDS and 100 mM NaHCO_3_ by two successive incubation at 55°C for 15 min. For the input control sample, 2% of fragmented chromatin was diluted with elution buffer, and treated as ChIP samples for subsequent steps. NaCl and RNaseA (Sigma-Aldrich) were added to input control and ChIP eluate to a final concentration of 200 mM and 100 *μ*g/mL, respectively, and incubated at 65 °C for 16 hours. 100 *μ*g/mL Proteinase K (Qiagen) was added to the solution and incubated at 55 °C for 1 h. DNA was extracted with ChIP Clean and Concentrator Kit (Zymo) following the manufacturer’s instructions. DNA was quantified using a Qubit dsDNA HS Assay Kit (Life Technologies). For preparing libraries for next-generation sequencing, procedures involving end repair, 3′ end adenylation, ligation with barcoded adaptors, and PCR amplification were performed with a TruSeq DNA Sample Preparation Kit 200–400 bp were size selected using AMPure XP magnetic beads. Libraries were quantified on the NanoDrop 1000 Spectrophotometer. Libraries were pooled and sequenced on the Libraries were sequenced on Illumina NextSeq platform, using 75bp, paired-end reads.

ChIP for Smchd1 on autosomes (i.e. this protocol does not solubilise the heterochromatin of the Xi) was performed as above with modifications. Following lysis of 4 × 10^7^ female *Smchd1*^GFP/GFP^ and *Smchd1*^+/+^ NSCs, cells were dounced 25 times in a cold, tight dounce homogoniser. Nuclei were centrifuged for 1 minute at 12,000 rpm at 4°C, and washed in IP buffer. Cells were resuspended in 40 *μ*L 1 × MNase buffer (New England Biolabs) with 1 × BSA (New England Biolabs) for every million cells. Nuclei were incubated at 37°C for 5 minutes. 500 U MNase (New England Biolabs) was added per million cells, and nuclei were incubated at 37°C for 5 minutes. EDTA was added to a final concentration of 10 mM, and nuclei were incubated on ice for 10 minutes. Nuclei were centrifuged for 1 minute at 12,000 rpm at 4°C, resuspended in 130 *μ*L IP buffer, and subjected to sonication using the Covaris S220 sonicator (peak power: 125, duty factor: 10, cycle/burst: 200, duration: 15 s). The lysate was cleared by centrifugation for 10 minutes at 12,000g at 4°C. Smchd1 was then immunoprecipitated, and chromatin purified as above.

ChIP for histone proteins was performed in female X_129_^XistΔA^X_Castaneus_ *Smchd1*^del/del^ and *Smchd1*^del/fl^ NSCs, crosslinked as above. 1 × 10^6^ cells were used per ChIP. Nuclei were extracted by washing three times in nuclear isolation buffer [20mM Tris pH 8.0, 10 mM NaCl, 2 mM EDTA pH 8.0, 0.5% Igepal CA-630, 1X c0mplete protease inhibitor (Roche)] pelleted each time by centrifugation at 1500 rpm for 5 minutes at 4°C. Nuclei were resuspended in sonication buffer [20 mM Tris pH 7.5, 150 mM NaCl, 2 mM EDTA, 1 % Igepal CA-630, 0.3% Sodium dodecyl sulfate, 1X c0mplete protease inhibitor (Roche)] and sonicated using the Covaris S220 sonicator (peak power: 105, duty factor:20, cycle/burst: 200, duration: 750 s). Sample was cleared by centrifugation at 12 000 × g for 20 minutes, and 1 volume of dilution buffer [20 mM Tris-HCl pH 8.0, 150 mM NaCl, 2 mM EDTA, 1% Triton X-100, 1X c0mplete protease inhibitor (Roche)] was added to cleared chromatin. Sample was pre-cleared with 20 *μ*L of Dynabeads Protein G (Life Technologies) blocked with 0.1 % Bovine Serum Albumin (BSA) for 2 hours. BSA was added to pre-cleared chromatin to a final concentration of 0.1 %. 1 % chromatin was taken as input. Immunoprecipitaion was performed overnight at 4°C with rotation using 2 *μ*g antibody (anti-H3K27me3 antibody from Millipore 07-499, anti-H3K27ac antibody from Abcam Ab4729). Immunoprecipitatied samples were captured by incubation with 20 *μ*L Dynabeads Protein G (Life Technologies) blocked with 0.1 % BSA for 2 hours. Beads were then washed twice each in wash buffer 1 [20 mM Tris pH 8.0, 150 mM NaCl, 2 mM EDTA, 1% Triton X-100, 0.15% SDS], wash buffer 2 [20 mM Tris pH 8.0, 500 mM NaCl, 2 mM EDTA, 1% Triton X-100, 0.1% SDS], wash buffer 3 [20 mM Tris pH 8.0, 250 mM LiCl, 2 mM EDTA, 0.5% Igepal CA-630, 0.5% sodium deoxycholate], and TE buffer [10 mM Tris pH 7.5, 1 mM EDTA]. Crosslink reversal, elution, and library preparation was performed as above.

Mapping was performed using Bowtie2^30^ v0.12.9. Sequencing data were analysed with the aid of the Seqmonk v1.36.1 or v1.38.2 software. For Smchd1 on the Xi, and ChIP for the histones, samples were pooled in Seqmonk, and peaks were called using the MACS style caller within the Seqmonk package (settings for 10,000 bp fragments, p < 1 × 10−05) using the whole cell extract as the control. For the Smchd1 ChIP peaks used in the in situ HiC analysis, peaks were called using MACS2 with down sampling and a p < 0.001, and the WCE samples used as the control. Enrichment analysis for histone domains was performed by determining the percentage coverage over 1 Mb bins along the female X chromosome. Smchd1 domains were called using Enriched Domain Detector (EDD) software^31^ v1.1.18, using the following parameters: max_CI_value = 0.25, required_fraction_of_informative_bins = 0.98, p_hat_CI_method = agresti_coull, log_ratio_bin_size = 10.

Seqmonk browser tracks were produced by quantifying regions over 1 000 kb bins, sliding by 500 bp each time for histone marks, or 10,000kb sliding by 5kb each time for Smchd1, then normalised for library depth with Match normalisation, and the appropriate H3 or isotype control sample subtracted for each IP. ChIP-seq heatmaps over ATAC-seq peaks were generated using SeqMonk’s built in Heatmap function.

### ATAC-seq

ATACseq was performed exactly as described in^32^. ATAC-seq libraries were subject to to 75 bp, paired-end sequencing on the Illumina NextSeq platform. Reads were aligned to mm10 using Bowtie2 v2.2.5 with the parameter −X 2000^30^. Mitochondrial reads were removed using Samtools, and duplicates were removed with Picard^33^. MACS2 peak calling was performed using v2.1.0.20140616 the following parameters: ---bdg --nomodel --shift −75 --extsize 150 --qvalue 0.05^18^. ATAC-seq data were analysed using Seqmonk v1.36.1. Reads under MACS2 peaks were quantified, normalised for library size. Differential ATAC-seq peak analysis was performed using edgeR statistics in Seqmonk. The GREAT tool was used to assess differential ATAC-seq peaks in respect to TSSs^34^. Differential ATAC-seq peaks were compared with H3K4me3-enriched regions, H3K4me1 enriched regions, Ctcf binding sites, RNA polymerase II binding sites and predicted enhancer sites identified in E14.5 mouse brain in Shen *et al*.^35^. These genomic co-ordinates of these features was transferred from mm9 to mm10 for analysis. Hypergeometric probabilities were calculated using the phyper function in R.

### Reduced Representation Bisulphite Sequencing (RRBS)

DNA was isolated from X_129_^XistΔA^X_Castaneus_ *Smchd1*^del/del^, *Smchd1*^del/fl^, *Smchd1*^del/+^ and *Smchd1*^fl/+^ NSCs using a DNeasy Blood and Tissue Kit (Qiagen). RRBS libraries were made from 100 ng DNA using the Ovation RRBS Methyl-Seq System (Nugen) as per manufacturers’ instructions. Libraries were quantified using using a Qubit dsDNA Assay Kit (Life Technologies) for sequencing on the Illumina NextSeq platform^36^. Libraries were subject to 75 bp, paired-end sequencing. Sequencing reads were mapped to a bisulphite-converted version of the n-masked mm10 genome described above which was created using Bismark v0.13.0. Adapter trimming was performed with TrimGalore v0.4.0, and 7 5’ most bases and 2 3’ most bases were trimmed based on m-Bias results from Bismark^37^. Reads were then split into either *Mus musculus castaneus* or *Mus musculus domesticus* using SNPsplit. Methylation calls were made using the Bismark Methylation Extractor^38^. Analysis was then performed in Seqmonk v1.38.2. Only CpG sites covered by more than 10 reads were considered for analysis.

### Western Blotting

Cells were treated with trypsin-EDTA and washed twice in cold MTPBS. The following steps were all performed with rotation at 4°C for 10 minutes, followed by centrifugation at 13,000rpm at 4°C for 5 minutes. Cells were lysed with 500 *μ*L KALB lysis buffer [150 mM NaCl, 50mM Tris-HCl (pH 7.5), 1% (vol/vol) Triton X-100, 1 mM EDTA (pH 7.5)] supplemented with 1 X c0mplete protease inhibitor (Roche), 2 mM Na3VO4 and 1 mM PMSF, and supernatant was collected as whole cell extract.

Protein concentration was quantified using a Pierce BCA Protein Assay Kit (Thermo Fisher Scientific). Proteins were resolved by reducing SDS-PAGE on 4-12% Bis-Tris gels (Novex) in MES buffer (Life Technologies), and transferred to a PVDF membrane by wet transfer for 1 hour at 100V. Membranes were blocked in 5% (w/v) skim milk powder in 0.1% Tween-20/PBS for 1 hour at room temperature. The membrane was probed with a primary antibody in blocking solution overnight at 4°C or for 1 hour at room temperature, washed over 30 minutes with 0.1% Tween-20/MTPBS (6 changes), incubated with the appropriate secondary antibody conjugated to HRP in blocking solution for 1 hour, and rinsed as above. Antibody binding was visualized using the Luminata ECL system (Millipore) following the manufacturer’s instructions.

### Whole Mount Skeletal Staining

Whole mount skeletal staining of *Smchd1*^MommeD1/MommeD1^ *Smchd1*^MommeD1/+^ and *Smchd1*^+/+^ E17.5 embryos was performed as described^39^. Essentially, embryos were collected following euthanisation of timed pregnant dams and washed in 1 × PBS. Eyes, skin, internal organs and adipose tissue were removed, and embryos were fixed in 95% (v/v) EtOH overnight at room temperature. Embryos were then moved into acetone overnight at room temperature. Cartilage was stained by incubating the sample in Alcian blue solution [0.03% w/v Alcian Blue 8GX, 80% (v/v) EtOH, 20 % (v/v) glacial acetic acid] overnight at room temperature. Embryos were destained by washing twice in 70% (v/v) EtOH followed by overnight incubation in 95% (v/v) EtOH. Tissue was pre-cleared by incubation in 1% (w/v) KOH for 1 hour at room temperature. Bones were stained by incubation in Alizarin red solution [0.005 % (w/v) Alizarin red in 1 % (w/v) KOH] for 4 hours at room temperature. Embryos were destained in 50% glycerol:50 % (1 %) KOH at room temperature until cleared, and stored in 100% glycerol until imaged on a ZEISS SV11 Stereomicroscope.

### Antibodies

For Western blotting for Smchd1, we used an in house produced rat monoclonal antibody raised against the Smchd1 ATPase domain, used at 1:1000 dilution. Anti-Actin antibody for Western blot was from Santa Cruz (H1015), used at 1:5000 dilution. For Smchd1-GFP ChIP, we used Life a Technologies antibody (A111-22), using 10 *μ*g antibody per ChIP. For H3K27me3 ChIP, we used a Millipore antibody (07-449), 2 *μ*g per ChIP. For H3K27ac ChIP, we used an Abcam antibody (Ab4729), 2 *μ*g per ChIP. For H3K27me3 immunofluorescence, we used the same Millipore antibody (07-449) at 1:100 dilution. For H2AK119ub immunofluorescence, we used an antibody from Cell Signalling Technology (#8240) at a 1:100 dilution. For immunofluorescence, the secondary antibody used was anti-Rabbit-488 (Life Technologies, A21206), at 1:500 dilution.

**Supplementary Table 13.**
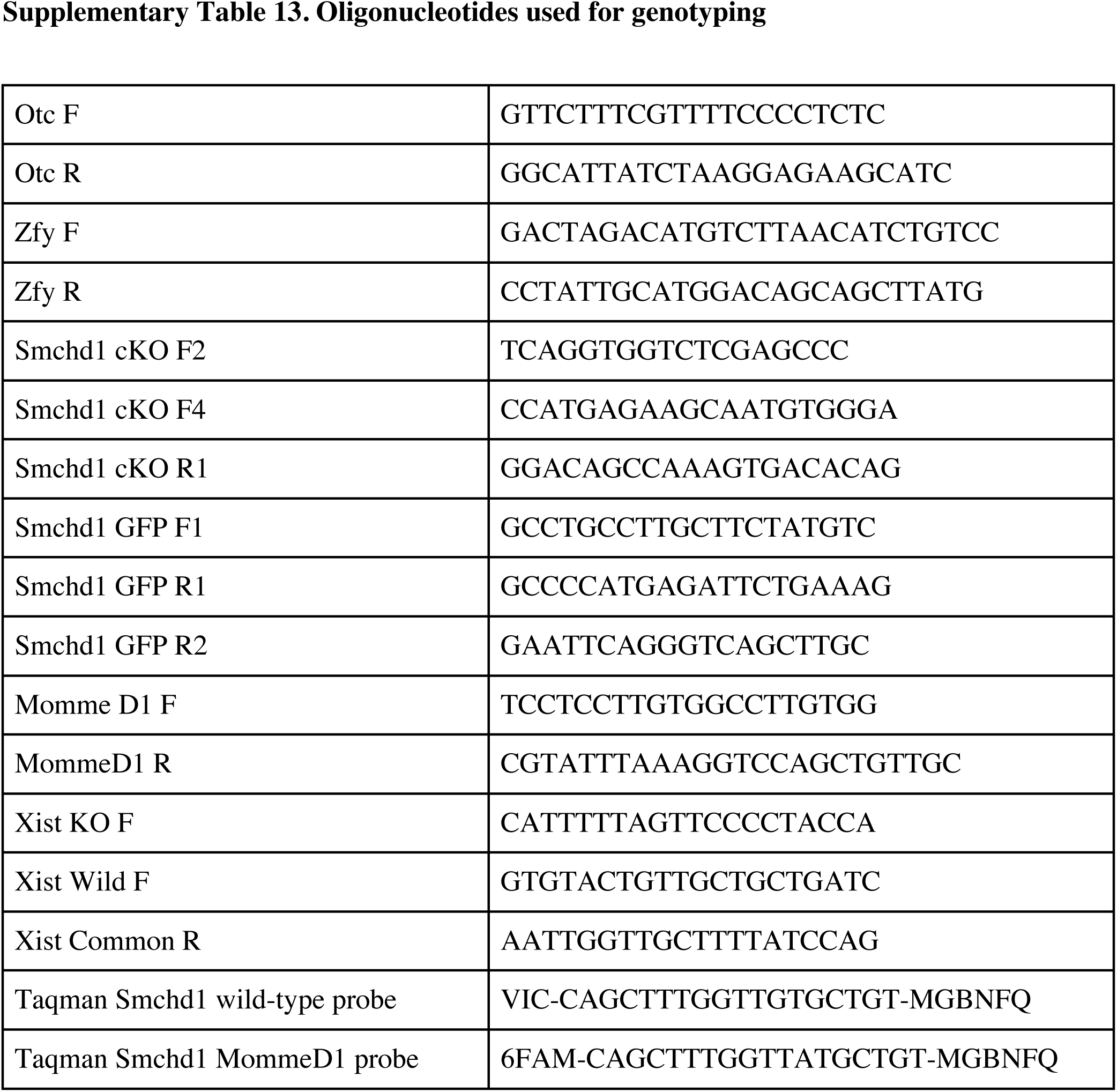
Oligonucleotides used for genotyping

